# Mechanism-driven screening of membrane-targeting and pore-forming antimicrobial peptides

**DOI:** 10.1101/2025.05.17.654650

**Authors:** Jiaxuan Li, Chenguang Yang, Ruihan Dong, Juan Francisco Bada Juarez, Lei Wang, Maximilian Emanuel Wettstein, Dali Wang, Chan Cao, Ying Lu, Chen Song

## Abstract

The rise of antibiotic resistance has generated an urgent demand for the discovery of new antimicrobial peptides (AMPs), prompting the development of various screening strategies. However, the specific function mechanisms of AMPs are often overlooked during the screening and optimization processes. In this study, we introduce a mechanism-driven screening approach that employs machine learning-based computational models to identify peptide sequences that target membranes and form pores. This approach explicitly considers critical factors such as structural features, membrane ainity, and the ability of peptides to oligomerize. Our method was applied to the metaproteomes of poison frogs, African clawed frogs, and human skin, followed by experimental validation. Seven peptides were successfully screened, each demonstrating antimicrobial activity with minimal hemolysis and cytotoxicity. These peptides exhibited membrane disruption capabilities in liposome leakage assays, with three showing broad-spectrum antimicrobial activity. Furthermore, single-molecule experiments indicated that these peptides can oligomerize on membranes, while electrophysiological measurements detected pore formation, confirming the effectiveness of our screening strategy. Therefore, our screening approach can effectively identify AMP sequences that act through membrane-targeting and pore-forming mechanisms, offering a promising, mechanism-driven strategy for the discovery of new antimicrobial agents to combat antibiotic resistance.

## 1 Introduction

Peptides that self-assemble into transmembrane pores can function as ion channels, ^1^ biosensors, and antimicrobial agents, ^2^ thereby demonstrating wide-ranging biomedical applications. ^3^ These peptides oligomerize and insert into the hydrophobic core of lipid bilayers, forming barrel-shaped pores that facilitate membrane permeation and lead to leakage of cytoplasmic contents. ^4^ Due to their membrane-disrupting characteristics, some pore-forming peptides favor rapid inhibition of a broad spectrum of bacteria. ^5^ These antimicrobial peptides (AMPs) are less likely to induce resistance, distinguishing them from conventional antibiotics. ^6^ Therefore, they represent ideal drug candidates with the potential to address the global crisis of drugresistant infections.

Designing novel pore-forming AMPs is challenging due to the limited amount of knowledge about their sequence and function mechanism. Only a few natural AMPs, such as melittin,^7^ alamethicin, ^8^ dermcidin,^9^ have been validated with a clear pore-forming mechanism. Therefore, a common approach is to use a template-based rational design, which builds a library of peptide variants with empirical substitutions and measures their activity through leakage experiments.^10–12^ Pore-forming peptides can also be engineered from truncated transmembrane domains of transporters, such as cWza ^13^ and DpPorA. ^14^ Meanwhile, molecular dynamics simulations provide computational insights on peptide assembly, pore formation,^15^ and pore stabilization,^16^ which can guide modifications to peptide sequence compositions and the design of pore-forming AMPs. ^17^ However, a significant limitation of rational design is its time-consuming nature, which hinders the exploration of diverse sequence candidates.

With the rapid advances in machine learning methodologies, computational mining has accelerated the discovery of new AMPs. Since AMPs are widely present in living organisms, researchers have successfully identified AMPs encoded by small open reading frames (smORFs) across numerous microbiomes. ^18–20^ Certain protein fragments can also act as AMPs, making the proteomes of human and other extinct organisms valuable sources of encrypted AMPs. ^21–23^ The search space can be further expanded to include all hexapeptides by applying a mining procedure that sequentially incorporates filtering, classification, ranking, and regression steps. ^24^ These studies have used data-driven models to discriminate whether a peptide is potentially antimicrobial and have determined AMP mechanisms during the experimental validation stage, rather than considering the membrane-targeting potential of peptides when constructing the computational pipeline. To the best of our knowledge, there are currently no mechanism-driven screening methods that focus on the discovery of pore-forming AMPs.

In this study, we established a machine learning-assisted and mechanism-driven strategy to identify new pore-forming AMPs, as shown in Fig. 1. First, we trained a support vector machine (SVM) predictor to screen membrane-active AMP candidates from given protein sequences, such as those from human and frog metaproteomes. Subsequently, the membrane contact probability (MCP), ^25^ helical propensity, and antiparallel interactions of these peptides were predicted to aid in ranking and selecting candidate peptides. In our design, these selected peptides were considered more likely to oligomerize on bacterial membranes and consequently more likely to form pores. Through *in vitro* validation, we found that seven out of 18 selected peptides exhibited antimicrobial activity, with the most effective exhibiting a minimum inhibitory concentration (MIC) of 2 µg/mL. These peptides also showed minimal cytotoxicity and hemolysis, and three of them showed broad-spectrum antimicrobial activity against Gram-positive and Gram-negative bacteria. Single-molecule photobleaching verified that these AMPs formed dimers, trimers, and tetramers upon interaction with lipid bilayers. All of the screened AMPs were able to induce membrane leakage, and three displayed pore-forming activity. Therefore, we have established a new screening pipeline designed to identify membrane-interacting and pore-forming antimicrobial peptides (AMPs). We believe that this machine learning-based and mechanism-driven strategy will facilitate the discovery of new antibiotic agents.

**Figure 1:**
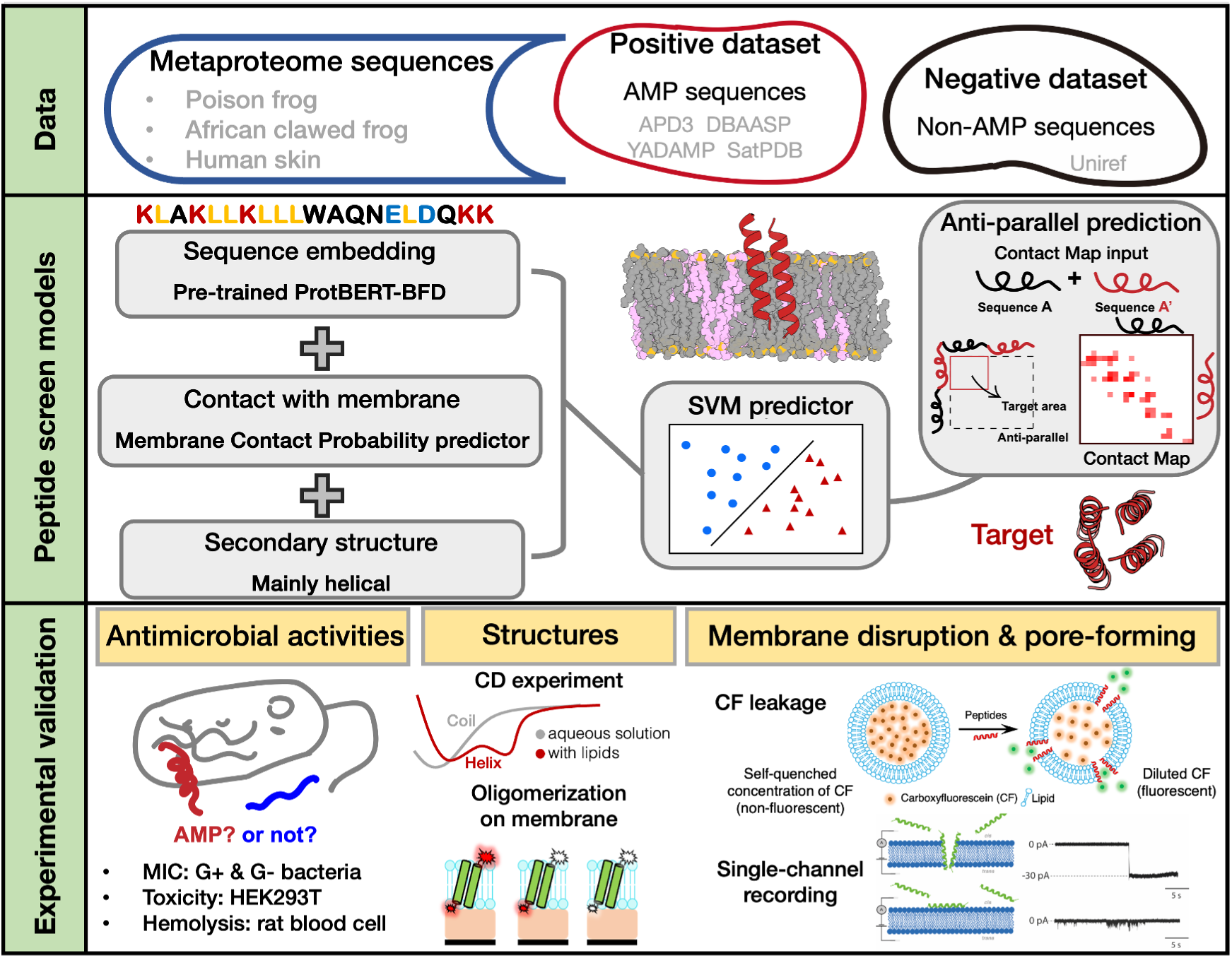
Overview of this work: Antimicrobial peptide candidates were identified using a Support Vector Machine (SVM) classifier enhanced with the ProtBERT-BFD protein language model. The screening process evaluated potential metaproteome sequences for membrane-targeting and pore-forming properties. Our classification system incorporated membrane contact probabilities and secondary structure predictions to identify peptides with strong membrane penetration potential. We prioritized sequences showing anti-parallel configurations, which are associated with membrane perforation activity. The selected peptides underwent comprehensive experimental validation to assess their antimicrobial effectiveness, structural properties, and membrane disruption mechanisms, with particular focus on their poreformation capabilities.

## 2 Results

### 2.1 Computational models for screening membrane-targeting and pore-forming AMPs

We developed a Support Vector Machine (SVM) peptide classifier using a pre-trained protein sequence encoder to identify sequences with antimicrobial properties. We chose the ProtBERT-BFD model,^26^ known for its high performance in encoding protein information, to generate feature vectors from input sequences. This approach takes advantage of the power of large language models, which have shown remarkable abilities to predict protein properties and functions without supervision. ^26,27^ Our classifier operates in the latent space created by the encoded sequences, as shown in Fig. 2A. To train the SVM model, we curated a positive dataset of AMPs from several reputable databases, including APD3, ^28^ DBAASP, ^29^ YADAMP, ^30^ and SATPdb. ^31^ To take advantage of the functional mechanisms and structural characteristics of membrane-associated AMPs, we predicted both the MCP ^25^ and the secondary structures for each sequence in this data set. In the positive data set, we included exclusively those peptides predicted to possess a helical structure with a high likelihood of membrane interaction, focusing on peptides ranging from 10 to 40 amino acids in length. We also created a negative dataset using UniRef, ^32^ selecting sequences without known antimicrobial properties. After training, we evaluated our model using 10-fold cross-validation with a Matthews Correlation Coeicient (MCC) of 0.93 and an accuracy that exceeded 0.98 (Table S1). We compared the performance of our model against other relevant methods, ^33–38^ and our model outperformed these approaches in four metrics on a benchmark dataset, ^38^ as shown in Table S2. These results demonstrate that our model is an effective antimicrobial peptide sequence classifier that is suitable for subsequent sequence screening.

**Figure 2:**
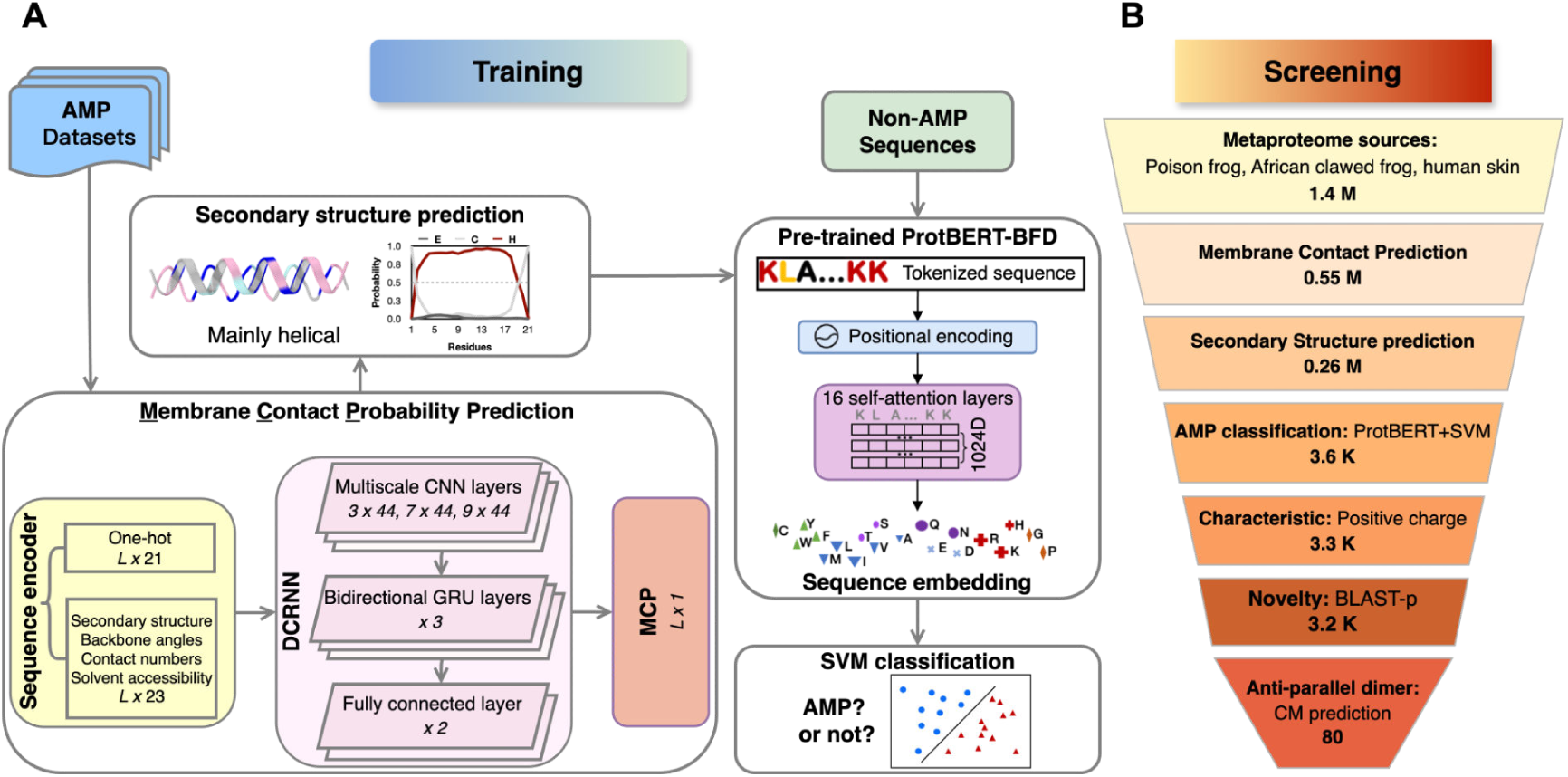
Mechanism-driven classification and screening process of antimicrobial peptides. (A) Training models for AMP classification with Surpport Vector Machine (SVM). Characteristics including membrane contact probability and secondary structure were predicted for the training dataset, and all sequences were encoded with the pre-trained ProtBERT-BFD model. Network architecture diagrams illustrate the model design. (B) Sequential screening and the corresponding number of sequences at each step. 80 AMP candidates were identified through computational analysis.

To further identify AMPs from a mechanistic perspective, we employed a multifaceted computational approach focusing on the membrane interaction and pore-forming potential of peptides. Our strategy used three key predictive steps, including the predictions of MCP, secondary structure, and oligomerization tendency. We used a deep learning-based MCP predictor to assess the likelihood that candidate peptides will interact with membranes. ^25^ This tool identifies amino acid residues of a given protein or peptide sequence that have a high probability of membrane contact, which is crucial for screening membrane-targeting sequences. Meanwhile, we focused on peptides predicted to have predominantly *α*-helical secondary structures, a common feature of transmembrane pore-forming AMPs. To evaluate the oligomerization potential of candidate peptides, we employed a ResNet-based contact map (CM) predictor, which has shown high precision in previous studies. ^25,39^ We applied this predictor to evaluate whether a candidate peptide can potentially form antiparallel dimers by examining the predicted contact maps. A diagonal contact pattern indicates a higher propensity for dimer or multimer formation. Fig. 1 (middle right) illustrates how the contact map’s target area corresponds to residue pairs in an antiparallel configuration. This approach allows us to identify peptides more likely to form oligomeric structures by antiparallel alignment, a key characteristic of some pore-forming AMPs that we studied previously. ^9,40^

We validated the effectiveness of our screening protocol by applying it to the metaproteome sequences of the poison frog, the African clawed frog, and the human skin in the Proteomics Identification Database. ^41^ We chose these metagenomic datasets as potential sources of AMP, as they are naturally rich in autoimmune and defense peptides, which aligns with recent studies that highlight the importance of targeted data selection in the discovery of new AMPs. ^19,21^ Detailed information on these models and datasets are available in the Materials and Methods section.

### 2.2 Screening criteria and preliminary evaluation

We designed a systematic set of criteria to screen for mechanism-interacting and pore-forming AMPs (Fig. 2B). We list the brief procedure here, and a more detailed description can be found in the Materials and Methods section.

1. Initial screening prioritized sequences with high MCP: An examination of known AMPs with membrane-interacting mechanisms showed that AMPs residues with predicted MCP values greater than 0.2 show a high probability of interacting with membranes (Details in Methods and Table S3). Metaproteome peptides with multiple residues that satisfy this criterion are selected for further screening.
2. Secondary structure screening: This analysis focused on peptides predicted to form *α*-helical secondary structure, a characteristic of membrane perforating AMPs (details in the Methods section and Table S3).
3. AMP classification with the SVM model: This screening step yielded a pass rate of 1.4% (3.6K out of 0.26M sequences) for peptides of 10-40 amino acids in length (Fig. 2B). The above process narrowed the candidates to 0.5%-1.7% of their initial sequences (detailed statistics in Table S4).
4. Characteristics: In the post-classification stage, we further checked the physicochemical properties of the candidate AMPs, particularly the positive charge distribution, to ensure conformity with typical AMP properties.
5. Sequence novelty: We evaluated sequence novelty through a comparison of BLAST-p with the APD3 database and used the E-value to select AMPs that did not exist in the AMP database previously.
6. Oligomerization potential: We analyzed the potential of the candidate peptides to form antiparallel dimers through contact map predictions of the spliced sequences. We use a new metric R_standard_ (Eq. (7)) to select candidate AMPs that show a diagonal contact map (see the method section for more details). 80 out of 3.2K sequences passed this step.

Fig. 2B presents the statistics of the sequence numbers during the screening process. This screening procedure resulted in the selection of 80 peptides, of which we selected 20 candidates with the highest perforated ability score F_score_. We define F_score_ as the product of the percentage of residues with high MCP in a certain peptide and the oligomerization capacity R_standard_ (Eq. (9)). This helped us identify the AMPs with a high probability of oligomerization on membranes. The sequences and predicted properties of all these 20 peptides are listed in Table S5.

We evaluated the primary physicochemical properties of these peptides, such as molecular weight and isoelectric point, against established AMPs within our positive datasets. The values of the screened peptides align with those of known AMPs (Fig. S1A). Furthermore, the amino acid composition of these peptides resembles that of known AMPs (Fig. S1B), supporting the validity of our screening process.

### 2.3 Antimicrobial activities of the candidate peptides

Following the above computational screening process, 20 candidate peptide sequences were subjected to experimental validation, among which, 18 sequences were successfully synthesized. These candidate AMPs were synthesized with a purity greater than 95% and prepared at a concentration of 1 mg/ml. We conducted minimum inhibitory concentration (MIC) experiments against 7 Gram-positive and 4 Gram-negative bacterial strains. As shown in Fig. 3, seven sequences demonstrated antimicrobial activity against at least one strain, and some of them showed broadspectrum antimicrobial activity. In particular, peptides 615, 623, and 1304 displayed significant inhibitory effects against Gram-positive and Gram-negative species, while peptide 58 exhibited an MIC value as low as 2 µg/ml against *S. hominis* ATCC 27842. Therefore, we obtained a 39% success rate (7 of 18) when translating from computational screening to experimental validation.

**Figure 3:**
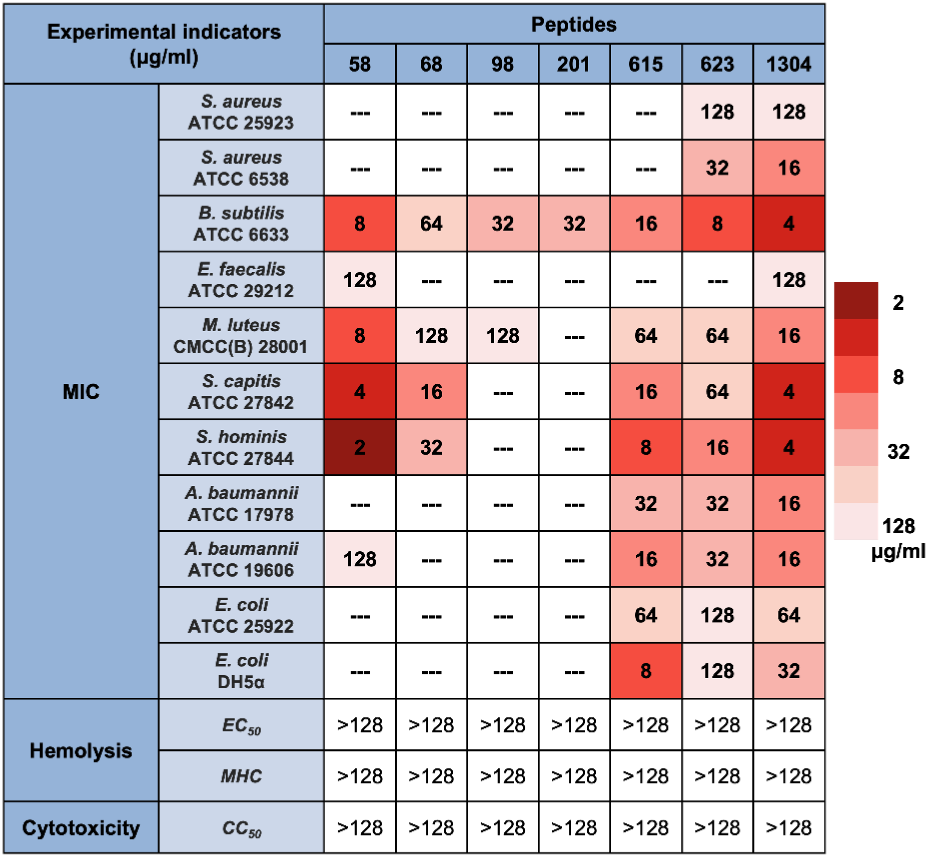
Antimicrobial eicacy and cell-damaging evaluation of selected seven peptide candidates. Experiments include: (1) Minimum Inhibitory Concentration (MIC) against seven Gram-positive and four Gram-negative bacterial strains. All peptides demonstrated potent antimicrobial activity and several exhibited broad-spectrum activity against multiple bacteria strains. (2) Hemolytic activity measured using rat red blood cells, with EC_50_ and MHC values exceeding 128 µg/ml. (3) Cytotoxicity analysis using HEK293T cells. All peptides exhibited low toxicity with a CC_50_ exceeding 128 µg/ml.

Fig. 4 illustrates the predicted characteristics of these seven peptides, indicating their abilities regarding membrane interaction and pore formation. These peptides consistently exhibited MCP values above 0.2 (Fig. 4A), helical structures (Fig. 4B), and clear antiparallel dimer formation patterns in the contact maps (Fig. 4C). Helical wheel analysis in Fig. 4D further demonstrated their amphipathic properties. Table 1 summarizes the characteristics of these seven AMPs, including their sequences, physicochemical properties, and predicted perforating score (F_score_). Most peptides are approximately 20 amino acids long and positively charged, with relatively high hydrophobicity, aligning with common AMP features. The novelty of these sequences was validated using BLAST-p with the APD3 database.

**Figure 4:**
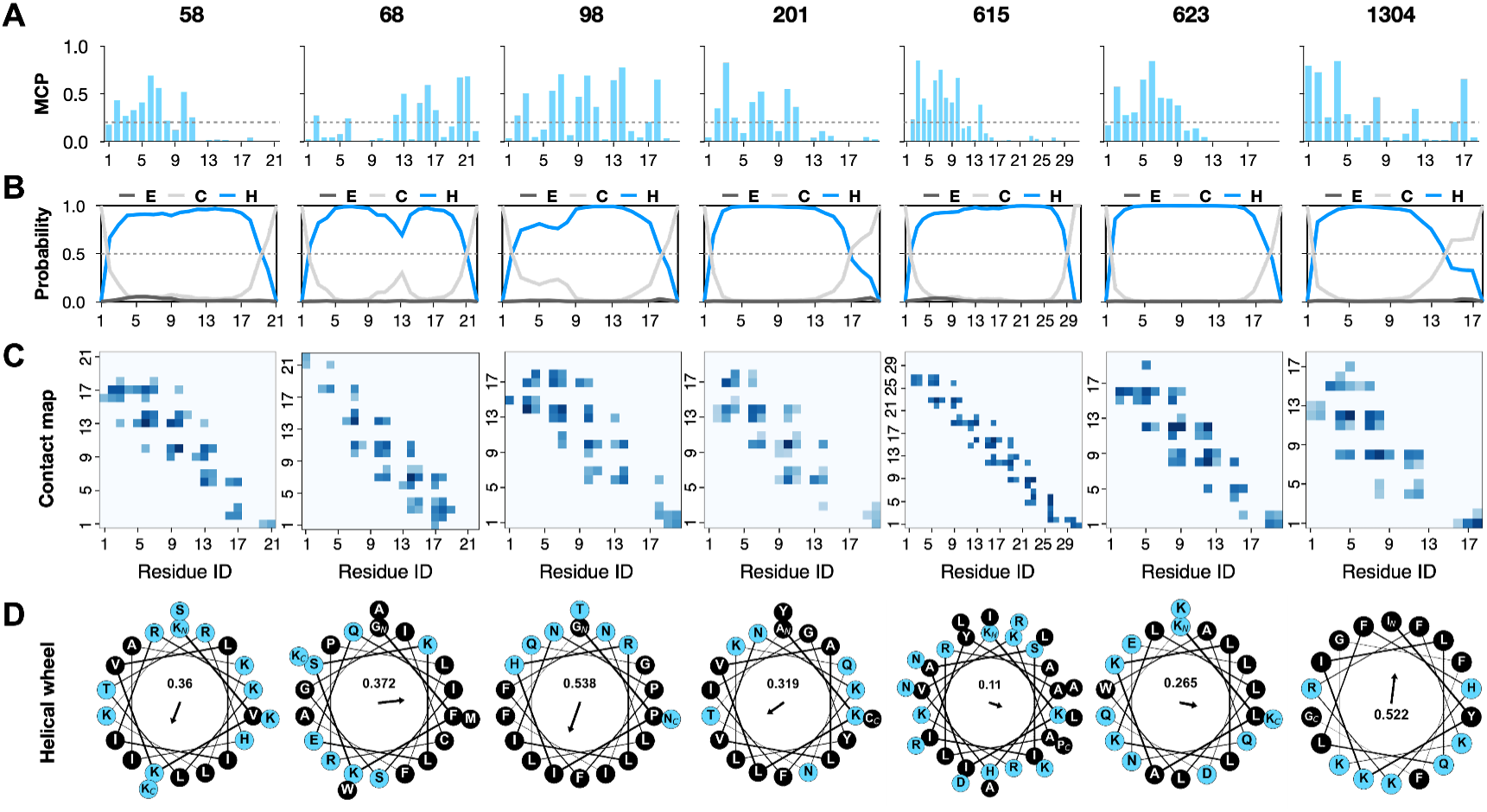
Characteristics of candidate sequences regarding membrane-targeting and pore-forming abilities. (A) Membrane contact probability predictions. Most of the residues in these seven peptides have predicted probabilities larger than 0.2, meaning they are likely to interact with membranes. (B) Secondary structure prediction results. Among three main secondary structures (E: *β*-sheet; C: Coil; H: *α*-helix), all sequences are predicted to be mainly helical. (C) Contact map predictions of antiparallel dimerization. All sequences share common diagonal contact patterns with high linear regression coeicients. (D) Helical wheel plots of candidate peptides. These plots illustrate the amphipathicity of the peptides, which is quantified at the center of the wheel.

**Table 1:** Sequences, physicochemical properties, and the predicted perforated ability (F_score_) of 7 antimicrobial peptides.

| Peptide name | Sequence | Charge | Length | GRAVY | Hydrophilicity | $F_{\text{score}}$ | E-value |
| --- | --- | --- | --- | --- | --- | --- | --- |
| 98 | GPIHRLINPFFNLLQGIFTN | +2 | 20 | 0.35 | 0.30 | 0.396 | 5.5 |
| 68 | GFKSKLEQISGICRPLFAAMWK | +3 | 22 | 0.10 | 0.27 | 0.332 | 8.7 |
| 1304 | IYKILQLFHKRFKKGFFG | +6 | 18 | -0.06 | 0.39 | 0.328 | – |
| 58 | KVKVLIIRKLTRHIAKLKSKK | +10 | 21 | -0.28 | 0.48 | 0.319 | 9.3 |
| 623 | KLAKLLKLLLWAQNELDQKK | +3 | 20 | -0.39 | 0.50 | 0.312 | – |
| 615 | KKIASIIYAHVKALRARKILDNLKRLAANRP | +9 | 31 | -0.22 | 0.42 | 0.292 | 7.1 |
| 201 | AKLVALVNKFIGYLKQNTYC | +3 | 20 | 0.37 | 0.30 | 0.264 | 7.8 |

Additionally, we performed cytotoxicity and hemolysis assays on these seven AMPs. As illustrated in the last three rows of Fig. 3, all the peptides exhibited outstanding biocompatibility, with toxicity and hemolytic activity well below conventional limits (CC_50_ > 50 µM, hemolysis < 10% at MIC). More details are provided in Table S6. At the maximum concentration tested, which was 128 µg/ml, all peptides showed excellent performance in key metrics, such as the half-maximal cytotoxic concentration (CC_50_) for HEK293T cells, the half-maximal effective concentration (EC_50_) for rat blood cells, and the minimum hemolytic concentration (MHC) defined as 10% hemolysis. In the cytotoxicity assay, the growth inhibition rates of seven AMPs were significantly lower than those for the Amphotericin B antibiotic used in clinical settings (see Table S7).

The peptides exhibit promising therapeutic potential, especially due to their demonstrated antimicrobial activity, along with low levels of cytotoxicity and hemolytic activity. However, it remains unclear whether their antimicrobial mechanisms align with our design, which will be discussed subsequently.

### 2.4 Experimental verification of membrane-targeting and pore-forming abilities of screened AMPs

We used single-molecule fluorescence photobleaching imaging to determine the stoichiometry of subunits in these peptides on supported lipid bilayers composed of POPE/POPG (3:1). ^42^ Based on the characteristics of the sequence of peptides, we adopted two labeling strategies for the seven selected AMP candidates. Peptides 68 and 201 that contain a single cysteine were labeled with Cyanine3-maleimide (Cy3) dye, which specifically reacts with the thiol groups on the cysteine residue of the peptides. Peptides 58, 98, 615, 623, and 1304, do not possess any cysteine, and they were labeled by an N-Hydroxy succinimide(NHS)-Rhodamine dye, in which the NHS esters react directly with the N-terminus of the peptides under acidic conditions (pH 4.5–6.5) to form amide bonds. ^43^ To avoid multiple labeling of NHS-Rho, peptides and fluorophores were incubated with a molar ratio of 1:1. The labeled peptides were continuously excited with low laser power (5 to 10 mW) to ensure slow photobleaching, minimizing the possibility of synchronous photobleaching of multiple fluorophores. The results of liposome leakage experiments showed that the fluorophores labeling on the peptides did not alter their membrane permeability, with the exception of peptide 68 (Fig. S2). We, therefore, excluded the single-molecule fluorescence data for the peptide 68.

TIRF microscopy images show that the densities of peptides 615, 623, and 1304 onthe lipid bilayer reached ∼0.2 particles/µm^2^ at low concentration (∼0.5 nM) while the density of peptide 58 decreased significantly to ∼0.1 particles/µm^2^ (Fig. 5A and S3). Moreover, the density of peptides 98 and 201 even decreased to < 0.05 particles/µm^2^(Fig. 5A and S3). This implied that peptides 615, 623 and 1304 have a comparable and strong capacity to absorb on the lipid bilayer. Compared to the three previously mentioned peptides, we observed that peptide 58 only possess half the adsorption capacity, while the adsorption of peptides 98 and 201 on the lipid bilayer is even lower. Additionally, we imaged peptide 68 by TIRF and observed an abnormal higherorder aggregation on the supported lipid bilayer, which is consistent with the result of liposome permeability measurement (Fig. 5A and S2).

**Figure 5:**
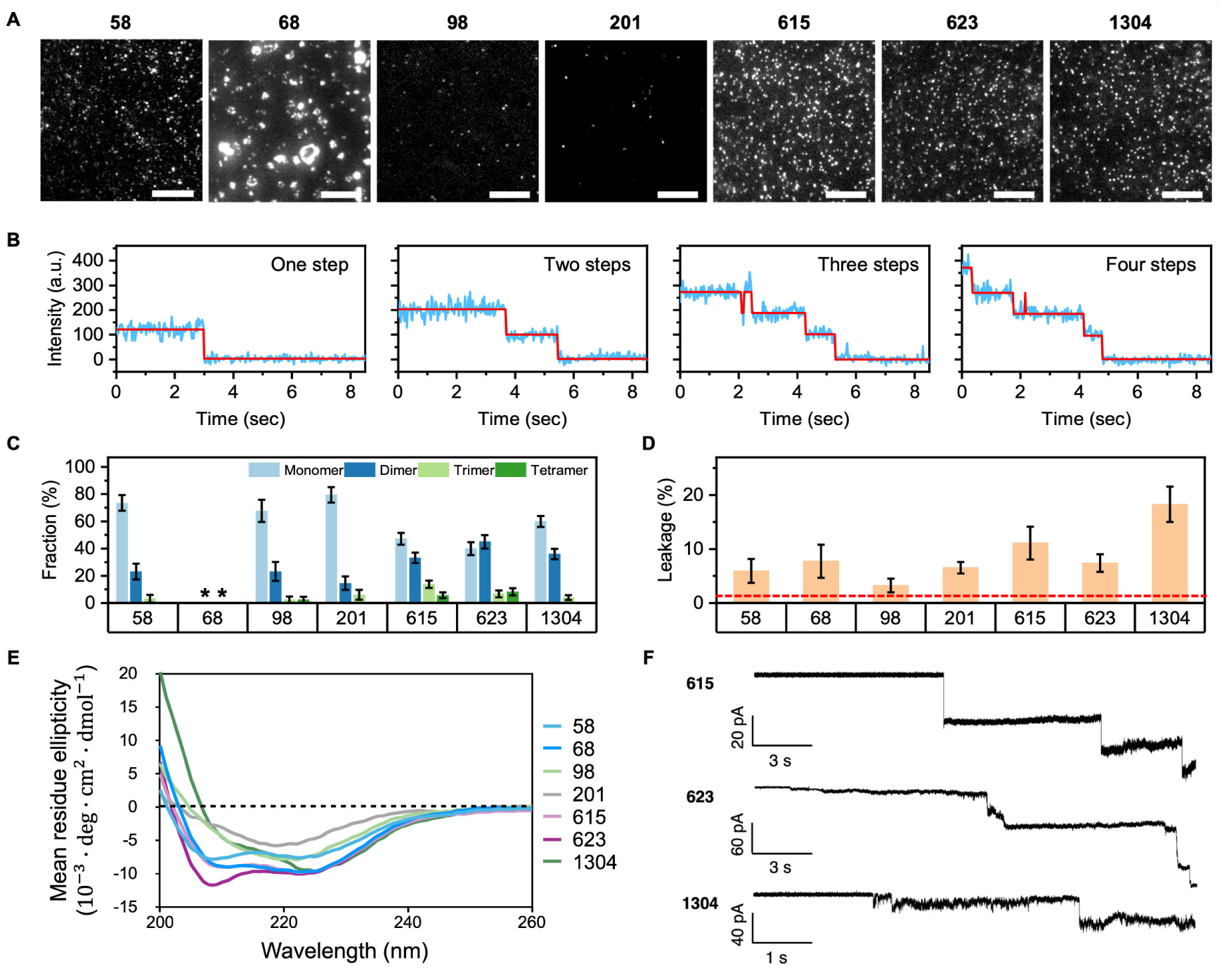
Experimental validation, correlating peptide oligomerization states with membrane disruption and pore activity. (A) The single-molecule TIRF image of fluorophores labeled peptide. Scale bar 10 µm. (B) Representative intensity traces (intensity trace, blue; idealized fitting curves, red) with time delivers multi-step photobleaching signals corresponding to monomer, dimer, trimer, and tetramer, respectively. (C) Histogram of photobleaching step numbers for fluorescent labeled pep-tides analyzed. Asterisks represent the data of peptide 68 (not further analyzed). Data are presented as mean ± SD. Error bars correspond to the bootstrapped standard deviation. (D) Leakage of carboxyfluorescein (CF) from liposomes induced by peptides. Histograms represent the leakage of CF from liposomes after the addition of peptide and are shown as a percentage of that induced by the addition of detergent (0.2% Triton X-100). Red dash lines indicate the background leakage from the liposomes incubated with an equivalent concentration of buffer alone, to control for the effect of buffer. Data are presented as mean ± SEM (n = 3 independent measurements). (E) Circular Dichroism (CD) experimental results with liposomes, showing the helical structure of most peptides. (F) Single-channel bilayer recordings of pore-forming signals, showing characteristic steps. The scale bar represents the current intensity and time scale respectively.

The fluorescence intensity trace of peptides on the lipid bilayer mainly exhibited one, two, three, and four photobleaching steps (Fig. 5B), indicating that peptides oligomerize on the lipid bilayer. The distribution indicates that all of the tested peptides can form dimers or even higher-order oligomers, which aligns with our design. Although monomers remain the predominant form for peptides 58, 98, and 201, pep-tides 615, 623, and 1304 exhibit a pronounced tendency to oligomerize, with more than 30% forming dimers (Fig. 5C). Interestingly, the liposome leakage measurements showed that peptides 615, 623, and 1304 have a stronger liposome leakage eiciency than peptides 58, 98, and 201 (Fig. 5D), which aligns with the proposed activity mechanism, wherein peptide oligomerization enhances membrane disruption and pore formation.

Importantly, the oligomers of the peptides were stable in the lipid bilayer, as the removal of the free peptides in solution did not significantly reduce the fraction of peptides as dimers (Fig. S4). In addition, single-molecule surface-induced fluorescence attenuation (smSIFA) was used to investigate the topology of peptide 201 on the membrane. ^44,45^ The findings revealed that peptide 201 assembles as an antiparallel dimer (Fig. S5), which is also consistent with our design. We were unable to conduct similar experiments for the other peptides due to the exclusive feature of a C-terminal Cy3 fluorophore in peptide 201, an essential requirement for precise membrane topology determination through this fluorescence-based methodology.. CD spectroscopy of the seven candidate AMPs with DOPE:DOPG (3:1) liposomes confirmed the predicted secondary structures during our screening procedure. As shown in Fig. 5E, peptides 58, 68, 615, and 623 displayed characteristic *α*-helical pattern, while peptides 98, 201, and 1304 showed a lower helical content, probably due to the existence of a flexible terminal coil., but these results align well with our predictions (Fig. 4B).

We also performed single-channel bilayer experiments to determine if the peptides were able to form pores or merely disrupt the membrane following the carpet model while in contact with it. We observed clear step transitions of different current amplitudes for the cases of peptides 615, 623, and 1304 (Fig. 5F and S6) indicative of pore formation. However, for peptide 68, we only observed a continuous increase in current over time (Fig. S6), suggesting a gradual disruption of the membrane without pore formation. For peptides 58, 98, and 201, no clear sign of pore-forming signals could be found. Interestingly, we only observed pore formation or peptide-membrane interaction only when PE:PG lipids (simplified model of bacterial membrane) were used, and no interaction was observed with pure PC lipids for several peptides. This lipid selectivity correlates well with our MIC, cytotoxicity, and hemolysis data (Fig. 3), where the peptides show only activity against bacteria (rich in PE and PG lipids), but not against HEK293T and rat red blood cells that are rich in PC lipids in the outer membrane.

## 3 Discussions

In this study, we developed a method that uses machine learning-based techniques to screen novel membrane-targeting and pore-forming AMPs. Our approach successfully identified seven effective antimicrobial peptides from metaproteomes. Subsequent experimental validation confirmed that these peptides can inhibit the growth of multiple strains of bacteria while exhibiting a low level of cytotoxicity. More importantly, they can bind and disrupt bacterial membranes, and three of them exhibited strong membrane ainity and oligomerization capabilities, as well as evidence of pore formation.

Unlike other deep learning-based AMP design methods that generally aim to increase the success rate, ^46–48^ our focus was on mechanism-driven screening. We would like to emphasize that while our approach may not achieve the highest success rate, it is capable of identifying AMPs that can form pores in bacterial membranes, a significant improvement in mechanism-driven AMP screening. Recently, Liu et al. conducted an interesting study in which they screened and evaluated AMPs self-assembly on bacterial membranes, yielding notably positive results. ^49^ Here, we introduce a more focused and challenging screening strategy specifically aimed at pore-forming AMPs. Consequently, the machine learning pipeline developed in this paper demonstrates that high-throughput screening of AMPs, targeting their precise functional mechanisms through a mechanism-driven approach, is now achievable. Investigating through *in vitro* experiments the peptide-membrane interaction, peptide oligomerization, and pore formation also provided insight into the antimicrobial activity of AMPs. We observed the formation of helical oligomers, including dimers, trimers, and tetramers, which are the fundamental components in the formation of transmembrane pores, which resemble barrel-stave structures. The barrel-stave model is one of the most important functional mechanisms of AMPs. ^7^ Our single-molecule experiments confirmed that peptides with larger membrane leakage and higher oligomeric states are those that effectively form pores on membrane (615, 623, 1304), as shown in Fig. 5 and S6. More interestingly, from Fig. 3, we can also infer that peptides 615, 623, and 1304 showed the broadest-spectrum antimicrobial activities. Thus, we hypothesize that pore-forming AMPs targeting bacterial membranes likely represent the most broad-spectrum candidates that deserve more thorough exploration.

Our approach, which combines computational prediction, experimental validation, and mechanistic insight, not only validates the effectiveness of our methodology but also paves the way for future research on the discovery and design of pore-forming AMPs. As discussed above, pore-forming AMPs carry great potential as novel antimicrobial agents, addressing the growing concern on antibiotic resistance. Meanwhile, the membrane-perforating mechanism of these peptides suggests potential applications beyond antimicrobial therapy, such as in anti-biofilm formation, cancer treatment, and membrane penetration for drug delivery.

In conclusion, our mechanism-driven and machine learning-based approach offers a promising strategy for the screening and design of membrane-pore-forming AMPs, with potentially far-reaching implications in medicine and biotechnology. Although further research is needed to fully realize its potential, this study represents a significant step toward antimicrobial drug discovery based on pore-forming AMPs.

## 4 Materials and Methods

### 4.1 Membrane contact probability (MCP) prediction

A previously developed membrane contact probability (MCP) predictor was used to assess the probability that a residue contact with membranes. ^25^ Since this model relied on evolutionary information derived from multiple sequence alignments (MSA), we used a modified MCP predictor without MSA input to handle AMPs with few homologous sequences. The input feature of a sequence (with length L) combines a one-hot vector (Lx21) with the output of SPIDER3-Single model, ^50^ which includes three-state secondary structure (Lx3), eight-state secondary structure (Lx8), accessible surface area (Lx1), main-chain angles (Lx8), half-sphere exposure (Lx2), and contact number (Lx1). The network architecture and other training settings are kept the same. For each input sequence, the MCP predictor can produce MCP values at the residue level.

### 4.2 Secondary structure prediction

We used the SPIDER3-Single model^50^ to predict the three-state secondary structures based solely on sequence data. This approach employed long short-term bidirectional recurrent neural networks with a one-hot vector as input, demonstrating greater accuracy compared to the evolutionary-based methods when dealing with proteins that have limited homologous sequences. The three-state secondary structure was defined using the DSSP program, consisting of α-helix (H), β-sheet (E), and coil (C) configurations. ^51^ SPIDER3-Single achieves a mean accuracy exceeding 70% across various databases, indicating its reliability in predicting three-state secondary structures. The cut-off value of H percentage was defined as 0.5 to identify α-helical AMPs.

### 4.3 Datasets

**Training dataset.** The dataset used for training SVM classifier involves a positive set of membrane-targeting AMPs and a negative set of non-AMPs. The positive set comprised 2141 natural AMP sequences sourced from established AMP databases, including APD3, ^28^ DBAASP, ^29^ YADAMP, ^30^ and SATPdb. ^31^ The lengths of these sequences ranged from 10 to 40 amino acids. Only α-helical peptides showing positive charges and having an MCP value of greater than 0.2 were kept. For the negative set, we selected non-AMPs from the UniRef database. ^32^ by filtering out sequences with keywords associated with antimicrobial activity, such as “antimicrobial”, “AMP”, “antifungal”, “anticancer”, “antibiotic”, “antiviral”, “toxic”, “defensive”, and “secretory”. Peptides with noncanonical amino acids were excluded as well. We maintained a positive-to-negative sequence ratio of 1:6 in the training dataset. 20% sequences were randomly selected as the validation set, keeping the remaining 80% for training.

**Benchmark dataset.** A balanced dataset comprising 4134 AMPs and 4134 non-AMPs was used to fairly compare our SVM model and other AMP predictors. ^38^ The splitting ratio of training and validation sets was also 8:2.

**Screening dataset.** The candidate metaproteome sequences were collected from the Proteomics Identification Database (PRIDE). ^41^ Three used metaproteomes were the poison frog (Project PXD002326 with 331,689 sequences), African clawed frog (Project PXD002326 with 999,997 sequences), and human skin (Project PXD020742 with 101,386 sequences). All sequences ranged from 10 to 40 amino acids in length.

### 4.4 SVM classifier

We encoded all input sequences using the ProtBERT-BFD model. ^26^ ThunderSVM was used to realize the SVM model with a radial basis function kernel. ^52^ The hyperparam-eters (’C’ and ‘gamma’) were optimized using grid search with cross-validations. The performance of the SVM classifier was evaluated by seven metrics, including sensitivity (Eq. (1)), specificity (Eq. (2)), accuracy (Eq. (3)), precision (Eq. (4)), Matthews Correlation Coeicient (MCC, Eq. (5)), F1 score (Eq. (6)) and area under the receiver operating characteristic curve (AUROC).

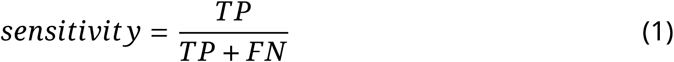

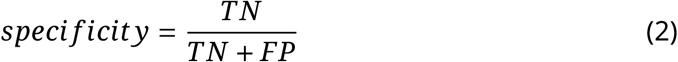

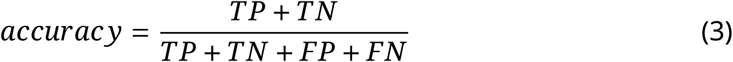

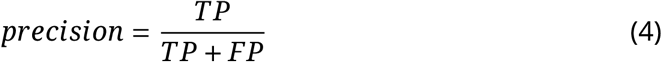

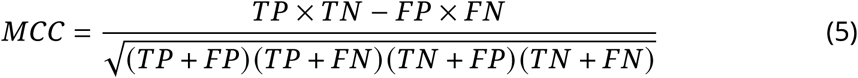

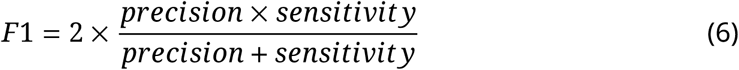

where TP is true positive, TN is true negative, FP is false positive, and FN is false negative.

### 4.5 Sequence analysis

The similarity and novelty of screened candidate AMPs were checked through BLAST-p^53^ in alignment with APD3 database. The cutoff of E-value was 10 for the sequence considered as novel AMP, as described in a previous study. ^54^ Physicochemical properties including charge and grand average of hydropathy (GRAVY) were calculated under standard criteria via the modlAMP package. ^55^ The GRAVY value is calculated by adding the hydropathy value of each residue and dividing by the length of the sequence. ^56^

### 4.6 Inter-chain contact map prediction and the perforated ability score

In line with the modified MCP predictor, we also used a MSA-free version of our MCP-incorporated contact map (CM) predictor^25,39^ to identify peptides that may oligomerize. Additionally, we employed the strategy of ComplexContact^57^ to extract interchain contacts between two peptide candidates and assess their potential of dimerization or even polymerization.

We predicted the contact map of the anti-parallel dimer through the ‘splice’ sequence of the twice-repeated peptide sequence. As shown in Fig. 1, the top left quarter of the whole CM is the target area of anti-parallel dimer prediction. The contact information between amino acids from the N to C terminal and inverse order were predicted. If the peptide was predicted to form a regular anti-parallel dimer, its CM resulted in a standard diagonal pattern, which could be quantified through R_standard_ (Eq. (7)). Here R^2^ is the coeicient of determination for linear regression in CM prediction and *L* is the length of the sequence.

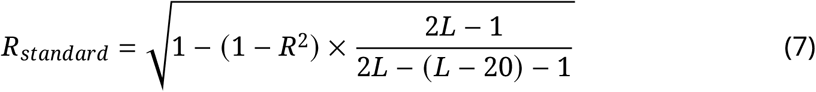

For the final evaluation of the screened peptides, we quantified their membrane contact ability (Eq. (8)) and oligomerization ability (Eq. (7)). F_score_ is the final score for AMP screening (Eq. (9)). Peptides with higher F_score_ are given priority for further experimental synthesis and validation (Table S5).

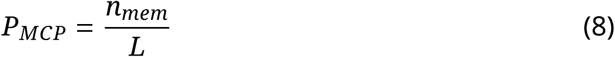

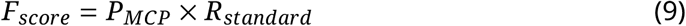

where *n_mem_* is the number of residues with predicted MCP larger than 0.2.

### 4.7 Peptides

Synthetic peptides 58, 68, 98, 201, 615, 623, 1304 were purchased from ChinaPeptides Co., Ltd. (purity > 95%, quality reports in Fig. S7).

### 4.8 Antimicrobial activity test

The minimal inhibitory concentration (MIC) of all screened peptides were determined by the microbroth dilution method. Bacterial strains included four Gram-negative strains (*Acinetobacter baumanii* ATCC 17978, *Acinetobacter baumanii* ATCC 19606, *Escherichia coli* ATCC 25922, *Escherichia coli* DH5α) and seven Gram-positive strains (*Staphylococcus aureus* ATCC 25923, *Staphylococcus aureus* ATCC 6538, *Bacillus subtilis* ATCC 6633, *Enterococcus faecalis* ATCC 29212, *Micrococcus luteus* CMCC(B) 28001, *Staphylococcus capitis* ATCC 27842, *Staphylococcus hominis* ATCC 27844). Bacteria were incubated overnight at 37^◦^C and diluted to 5 × 10^5^ cfu/mL. Then 99 µL bacterial suspension was added into each well of a 96-well plate. Peptides were dissolved in cation adjusted Mueller-Hinton (CAMHB) medium at an initial concentration of 1 mg/mL and then diluted in two-fold gradient concentrations, ranging from 0.125 to 128 µg/mL with 1 µL in each well. The plate was placed in incubator for 20 hours at 37^◦^C. The lowest concentration of peptide causing no observed bacteria grow was its MIC.

### 4.9 Hemolysis assay

The hemolysis was tested using rat red blood cells (RBC). Fresh rat blood were centrifuged at 500 g for 5 mins, then RBC precipitates were kept and resuspended with PBS (pH 7.4). RBC solutions were diluted and added into the 96-well plate at a density of 1 × 10^8^ cells per well. Peptides were diluted in PBS and added into the plate with final concentrations ranging from 0.125 to 128 µg/mL, with two-fold gradients. Triton X-100 (10 mg/mL) served as the positive control, resulting in 100% hemolysis. PBS was used as vehicle. After incubating for 1 hour, the plate was centrifuged at 500 g for 5 mins. The absorbance of supernatant was measured at 450 nm. The percentage of hemolysis was calculated as (OD_450, peptide_ - OD_450, PBS_) / (OD_450, positive_ - OD_450, PBS_). Half maximal hemolysis concentration (EC_50_) and lowest peptide concentration causing 10% hemolysis (MHC) values of each peptide were reported. EC_50_ was fitted with GraphPad Prism. Each test was conducted in triplicate.

### 4.10 Cytotoxicity test

Cytotoxicity was evaluated using the Alamar Blue assay on HEK293T cells. HEK293T cells were cultured in DMEM supplemented with 10% FBS and incubated at 37^◦^C with 5% CO_2_. Amphotericin B and staurosporine were used as controls. Peptide solutions were diluted in PBS (pH 7.4) with final concentrations ranging from 1 to 128 µg/mL. The cells were treated with peptides for 72 hours. After that, 10 µL Alamar Blue was added into each well. The plate was incubated for 4 hours under dark condition and detected the absorbance at 570 nm. The inhibition rate of cell growth was calculated via (OD_570, peptide_ - OD_570, PBS_) / (OD_570, staurosporine_ - OD_570, PBS_). The half maximal cytotoxic concentration (CC_50_) was fitted with GraphPad Prism. Each test was conducted in triplicate.

### 4.11 Preparation of supported lipid bilayers

All lipids were purchased from Avanti Polar Lipids and were dissolved in chloroform. Lipids with the following indicated compositions were mixed in a glass vial: 75% 1-palmitoyl-2-oleoyl-sn-glycero-3phosphoethanolamine (POPE), 25% 1-palmitoyl-2-oleoyl-sn-glycero-3-phospho-(1’-rac-glycerol) (POPG) (mol%). The lipid films were obtained by chloroform evaporation under a stream of nitrogen and then placed in vacuum pumps overnight to remove residual chloroform. The lipid films were resuspended to 2 mg/mL at room temperature with HEPES buffer (20 mM HEPES, 150 mM NaCl, pH 7.4) and sonicated until the suspension became transparent. 100 µL pre-prepared transparent vesicle solution was injected slowly into the imaging chamber and incubated at 37^◦^C overnight to form supported lipid bilayers. Then the remaining vesicles were washed away with HEPES buffer and the chamber used subsequently after the bilayer preparation.

### 4.12 Peptide-induced leakage of liposomes

To prepare 5(6)-carboxyfluorescein (CF)-encapsulated liposomes, lipid films composed of 75% POPE and 25% POPG were prepared as mentioned above. Then lipid films were hydrated in HEPES buffer containing 50 mM CF. A cycle of 10 freeze and thaw from liquid nitrogent to a 37^◦^C water bath were performed. Liposomes were generated by extruding the hydrated lipid films (21 passages.) through a 100 nm polycarbonate filter (Whatman) with a Mini-Extruder device (Avanti Polar Lipids). The non-encapsulated CF was then removed using a PD10 column (GE Healthcare). For the liposome leakage assay, 5 µg/mL lipid concentration of CF-encapsulated liposomes and 100 nM peptide proteins were incubated for 1h in HEPES buffer (20 mM HEPES, 150 mM NaCl, pH 7.4) at room temperature, and the emission fluorescence intensity (I) at 517 nm (excitation wavelength at 494 nm) recorded using a spectrophotometer (U3900H, Hitachi). The background fluorescence intensity I_0_ was monitored before the peptides were added. At the end of the incubation, the addition of 0.2% Triton X-100 was performed to measure the maximum fluorescence fluorescence intensity I_m_ corresponding to complete release of CF. The percentage of leakage was calculated as Leakage = [(I – I_0_) / (I_m_ – I_0_)] × 100%.

### 4.13 Fluorescence labeling

For fluorescence labeling of peptides, peptides were incubated with NHS-Rhodamine (NHS-Rho, Thermo Fisher) or sulfo-Cyanine3 maleimide (Cy3, Lumiprobe) at a 1:1 molar ratio at 4 ^◦^C for 12 h in the dark. To improve the yield of NHS-Rho labeling of the N-terminal amine, peptides 58, 98, 615, 623 and 1304 were labeled in the acidic HEPES buffer (20 mM HEPES, 150 mM NaCl, pH 5.4) because the N-terminal α-amine will be more nucleophilic than lysine ε-amine under acidic conditions. ^43^ In addition, peptides 68 and 201 were labeled with Cy3 in the neutral HEPES buffer (20 mM HEPES, 150 mM NaCl, pH 7.4). After labeling, the peptides were snap-frozen in liquid nitrogen and stored at –20 ^◦^C until needed.

### 4.14 Preparation of GO-BSA-supported lipid bilayer

The graphene oxide (GO) was produced by using the modified Hummer’s method and ultra-large GO flakes (supernatant) were collected through a multiple centrifugation-dispersion procedure. ^58^ The GO monolayer was deposited on the cleaned coverslips by using the Langmuir-Blodgett method. The coverslips and slides were sequentially pre-cleaned by acetone, methanol, and piranha solution, then rinsed with distilled water and dried out by nitrogen gas. Using the Langmuir-Blodgett equipment to deposit GO monolayer on the surface of the cleaned coverslips and then heat the deposited coverslips under vacuum at 85^◦^C for 2 hours to remove residual solution. Fluidic chambers with inlets and outlets were made by confining the GO-deposited coverslips to clean slides with double-sided tape (50 µm thickness, 3M). For GO-BSA-supported lipid bilayers, 1 mg/mL BSA dissolved in HEPES buffer was injected into the fluidic sample chamber and incubated for 30 minutes and then washed excess BSA solution with HEPES buffer. After preparation of BSA cushion, 100 µL pre-prepared transparent vesicles solution was injected slowly into the chamber and incubated at 37^◦^C overnight to form supported lipid bilayers. Then the extra vesicles were washed away by HEPES buffer. The chamber should be used as soon as possible after the bilayer preparation.

### 4.15 Single molecule photobleaching imaging and analysis

The single molecule photobleaching imaging was carried out using a TIRF microscope (Nikon, Ti2), equipped with a high numerical apertures oil immersion objective (Nikon, 100x, N.A. 1.49, oil immersion). A concentration of 0.25 nM peptides was added into imaging chamber and incubated for 10 min in HEPES buffer (20 mM HEPES, 150 mM NaCl, pH 7.4). Fluorophores were excited using a single-frequency 532-nm laser (Coherent, OBIS LS), and the emitted fluorescence signal was separated by a bandpass filter (Chroma, ET585/65m) and recorded by an EMCCD camera (Andor, IX897) with a frame interval of 100 milliseconds. The data was recorded as 16-bit movies in tiff format and further analyzed. Fluorophore coordinates were obtained using Particle Tracker, a plugin in ImageJ. The intensity trajectories of fluorophores were extracted based on the positions generated by Particle Tracker. The idealized photobleaching stepwise curves were determined using a custom Hidden Markov Model MATLAB script. ^59^

### 4.16 Circular dichroism experiments

1,2-dioleoyl-sn-glycero-3-phosphoethanolamine (DOPE) and 1,2-dioleoyl-sn-glycero-3-phospho-(1’-rac-glycerol) (DOPG) lipids were resuspended in chloroform and mixed at a molar ratio of 3:1. Chloroform was evaporated under a stream of nitrogen and the lipid film was hydrated with 600 µL buffer (20 mM Tris, 200 mM KCl, pH 7.4). After 10 freeze-thaw cycles, liposomes were extruded 20 times through a 0.2 µL pore-size filter.

The far-UV CD spectra of the different peptides with liposomes were acquired on a Chirascan V100 CD spectrometer with quartz cuvettes of 1 mm optical path in the buffer (20 mM Tris, 200 mM KCl, pH 7.4). The peptide concentration was set to 15 µM and the lipid concentration to 450 µM. The CD signal was measured from 200 nm to 260 nm at rate of 1 nm/s. Each spectrum presented here is an average of three spectra and data was not analyzed upon detector voltage saturation (below 200 nm). Data were smoothed in Prism 10 and the content in α-helix, β-sheet, and random coil was extracted from Dichroweb with the K2D algorithm and with BestSel.

### 4.17 Single-channel bilayer experiments

1,2-diphytanoyl-sn-glycero-3-phosphoethanolamine (DPhPE) and 1,2-diphytanoyl-sn-glycero-3-phospho-(1’-rac-glycerol) (DPhPG) powder were dissolved in octane (Sigma-Aldrich, Buchs SG, Switzerland) to a final concentration of 8 mg/mL and used in mixture as specified in the figure caption. Single-channel recording experiments were performed on an Orbit 16 TC instrument (Nanion). Phospholipid membranes were formed across a MECA 16 Recording Chip that contains 16 circular microcavities (100 µm diameter) in a highly inert polymer. Each cavity contains an individual integrated Ag/AgCl microelectrode and can record 16 artificial lipid bilayers in parallel. The buffer (20 mM HEPES, 200 mM KCl, pH 7.4), the concentration of peptides (2 µM), and the temperature (25^◦^C) were kept constant for all experiments. Membranes were formed and their capacitance was recorded. The traces were recorded in Elements Data Reader (Elements) and further analyzed by Clampfit (Molecular device). Data was collected at 20 kHz sampling rate. Results, fitting, and graphs were produced in GraphPad Prism 10.

## 5 Data and Code Availability

All datasets and codes will be available after peer review at https://github.com/ ComputBiophys/Pore-Forming_AMP_SVM.

## 7 Acknowledgement

The authors express their gratitude to Kai Kang and Hanyuan Sun for their insightful discussions. This work was supported by the National Key R&D Program of China (2024YFA0916800) and the Science Fund for Innovative Research Groups of the National Natural Science Foundation of China (T2321001). C.S. was supported in part by the Frontier Innovation Fund of Peking University Chengdu Academy for Advanced Interdisciplinary Biotechnologies. Part of the computation was performed on the computing platform of the Center for Life Sciences at Peking University. This research was also supported by the Swiss National Science Foundation (PR00P3_193090 to C.C.), Dementia Research Switzerland – Synapsis Foundation (2019-CDA02 and 2022-CDA03 to C.C.) and the Novartis Foundation for Medical-Biological Research.

## Supplementary Information

### 1 Supplementary Tables

**Table S1:** 10-fold cross-validations of our SVM predictor.

| Fold | Sensitivity | Specificity | Accuracy | Precision | MCC | F1 | AUROC |
| --- | --- | --- | --- | --- | --- | --- | --- |
| 1 | 0.9820 | 0.9918 | 0.9910 | 0.9213 | 0.9463 | 0.9507 | 0.9869 |
| 2 | 0.9671 | 0.9913 | 0.9894 | 0.9074 | 0.9311 | 0.9363 | 0.9792 |
| 3 | 0.9800 | 0.9925 | 0.9915 | 0.9188 | 0.9443 | 0.9484 | 0.9862 |
| 4 | 0.9607 | 0.9894 | 0.9867 | 0.9048 | 0.9250 | 0.9319 | 0.9751 |
| 5 | 0.9706 | 0.9907 | 0.9888 | 0.9116 | 0.9346 | 0.9402 | 0.9806 |
| 6 | 0.9627 | 0.9925 | 0.9899 | 0.9226 | 0.9370 | 0.9422 | 0.9776 |
| 7 | 0.9477 | 0.9936 | 0.9894 | 0.9368 | 0.9364 | 0.9422 | 0.9706 |
| 8 | 0.9481 | 0.9936 | 0.9899 | 0.9299 | 0.9335 | 0.9389 | 0.9708 |
| 9 | 0.9467 | 0.9907 | 0.9867 | 0.9091 | 0.9205 | 0.9275 | 0.9687 |
| 10 | 0.9551 | 0.9896 | 0.9867 | 0.8922 | 0.9160 | 0.9226 | 0.9724 |
| **average** | 0.9621 | 0.9916 | 0.9890 | 0.9155 | 0.9325 | 0.9381 | 0.9768 |
| **std** | 0.0123 | 0.0014 | 0.0017 | 0.0124 | 0.0092 | 0.0084 | 0.0061 |

**Table S2:** Benchmarking performances of various models using validation dataset. Best performances are shown in bold.

| Year | Method | Model | Sensitivity | Specificity | Accuracy | Precision | MCC | F1 | AUROC |
| --- | --- | --- | --- | --- | --- | --- | --- | --- | --- |
| 2018 | AMPScannerV2<sup>1</sup> | CNN+LSTM | 0.7860 | 0.8706 | 0.8283 | 0.8587 | 0.6590 | 0.8212 | 0.9010 |
| 2020 | MACREL<sup>2</sup> | RF | 0.7799 | 0.8670 | 0.8235 | 0.8543 | 0.6494 | 0.8160 | 0.8932 |
| 2021 | AmPEPpy<sup>3</sup> | RF | 0.7050 | 0.8948 | 0.7999 | 0.8701 | 0.6109 | 0.7738 | 0.8741 |
| 2022 | AMPLify<sup>4</sup> | LSTM+CNN | 0.7789 | **0.9033** | 0.8416 | **0.8897** | 0.6884 | 0.8307 | 0.9069 |
| 2022 | AMP-BERT<sup>5</sup> | BERT+MLP | 0.7993 | 0.8756 | 0.8374 | 0.8652 | 0.6767 | 0.8309 | 0.8374 |
| 2023 | Cao et al.<sup>6</sup> | BERT+MLP | 0.8186 | 0.8875 | 0.8531 | 0.8792 | 0.7079 | 0.8471 | **0.9141** |
| 2023 | AMPpred-MFA<sup>7</sup> | LSTM+CNN+Attention | 0.7545 | 0.8887 | 0.8216 | 0.8715 | 0.6492 | 0.8088 | 0.8942 |
|  | Ours | BERT+SVM | **0.8247** | 0.8948 | **0.8597** | 0.8869 | **0.7212** | **0.8546** | 0.8597 |

: second-best performances

**Table S3:**
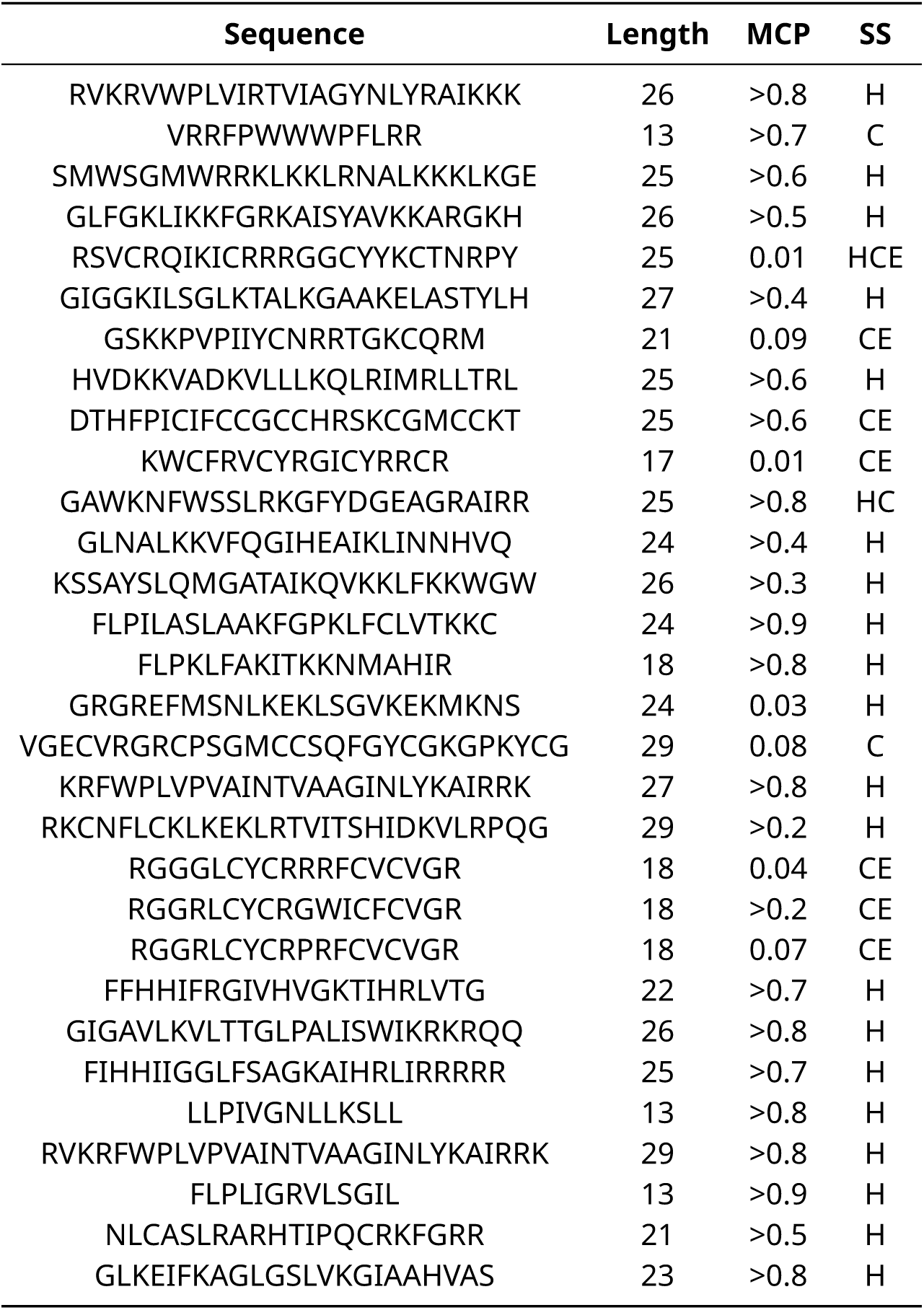
Benchmarking MCP and secondary structure (SS) prediction criteria with labeled sequences in DBAASP. H: α-helix, E: β-sheet, C: coil.

**Table S4:** Statistics of the sequence number in the screening process. SVM predictor and anti-parallel dimer CM prediction are rate-determining steps, where very few sequences could meet the expectations of mechanisms.

| Metaproteome source | Initial sequences | AMP classification |  |  | Post-classification |  |  |
| --- | --- | --- | --- | --- | --- | --- | --- |
|  |  | Pass MCP | Pass SS | Pass SVM | Positive charge | BLAST-p | Anti-parallel dimer (CM) |
| Poison frog | 331689 | 119013 | 51573 | 268 (0.5%) | 227 | 226 | 11 (4.9%) |
| African clawed frog | 999997 | 399781 | 189424 | 3250 (1.7%) | 2986 | 2956 | 67 (2.3%) |
| Human skin | 101386 | 36475 | 15968 | 75 (0.5%) | 56 | 56 | 2 (3.6%) |

**Table S5:**
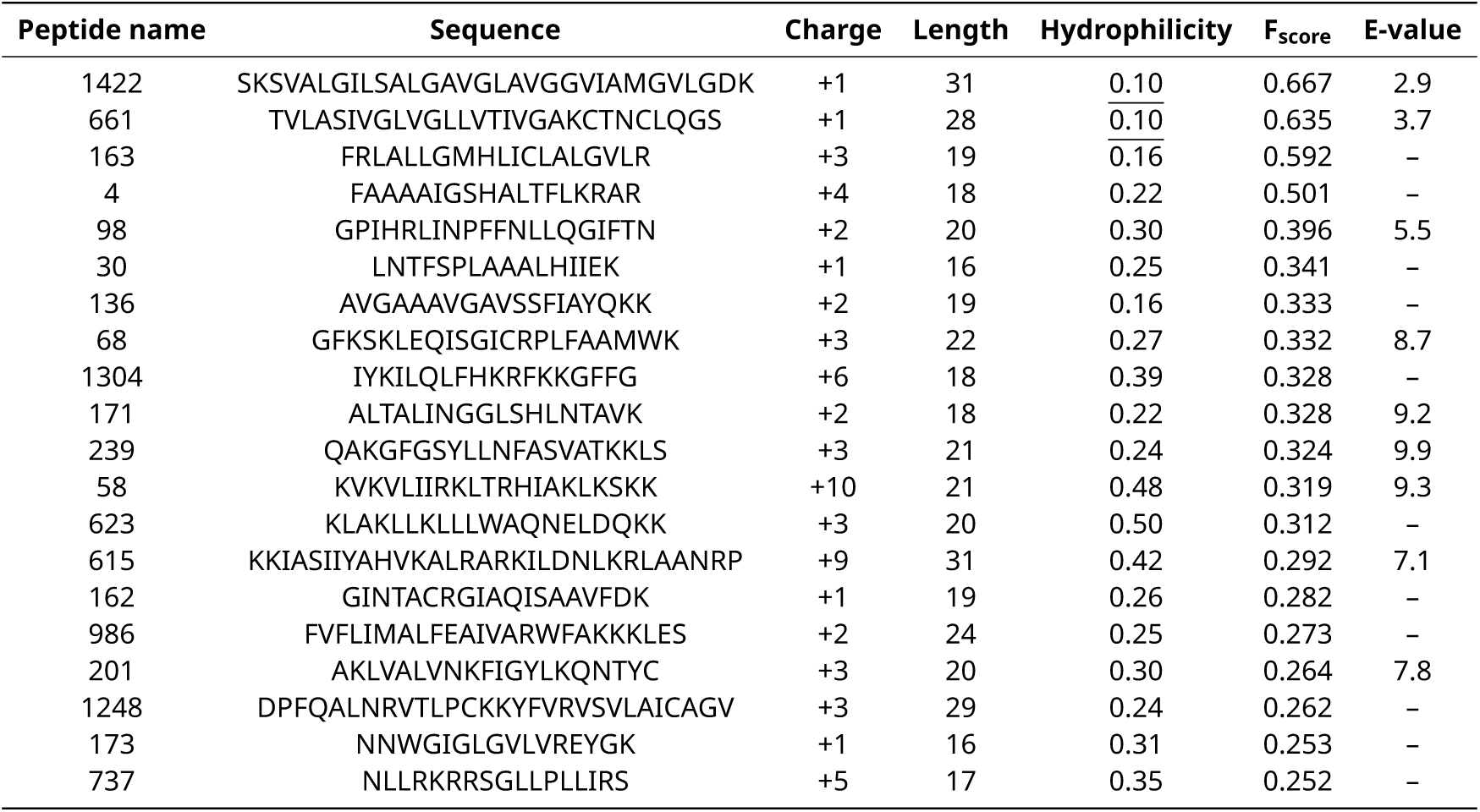
Sequences, physicochemical properties, and the predicted perforated ability (F_score_) of 20 selected antimicrobial peptides. Two sequences with underline are too hydrophobic for chemical synthesis.

**Table S6:** Hemolysis tests on rat blood cell (RBC). Both EC_50_ and MHC are larger than 128 µg/ml for all seven peptides. MHC: the lowest concentration of peptide that causes 10% hemolysis. Positive control : 10 mg/ml Triton X-100 + RBC with 100% hemolysis.

| Compounds | $EC_{50}$ ($\mu\text{g/ml}$) | MHC ($\mu\text{g/ml}$) |
| --- | --- | --- |
| 58 | >128 | >128 |
| 68 | >128 | >128 |
| 98 | >128 | >128 |
| 201 | >128 | >128 |
| 615 | >128 | >128 |
| 623 | >128 | >128 |
| 1304 | >128 | >128 |

**Table S7:** Cytotoxicity tests of seven peptides on HEK293T cells. CC_50_ is larger than 128 µg/ml for all screened peptides. Controls: Amphotericin B and Staurosporine.

| Compounds | HEK293T |
| --- | --- |
|  | CC<sub>50</sub> (µg/ml) |
| 58 | >128 |
| 68 | >128 |
| 98 | >128 |
| 201 | >128 |
| 615 | >128 |
| 623 | >128 |
| 1304 | >128 |
| Amphotericin B | 67.06 |
| Staurosporine | 0.26 |

### 2 Supplementary Figures

**Figure S1:**
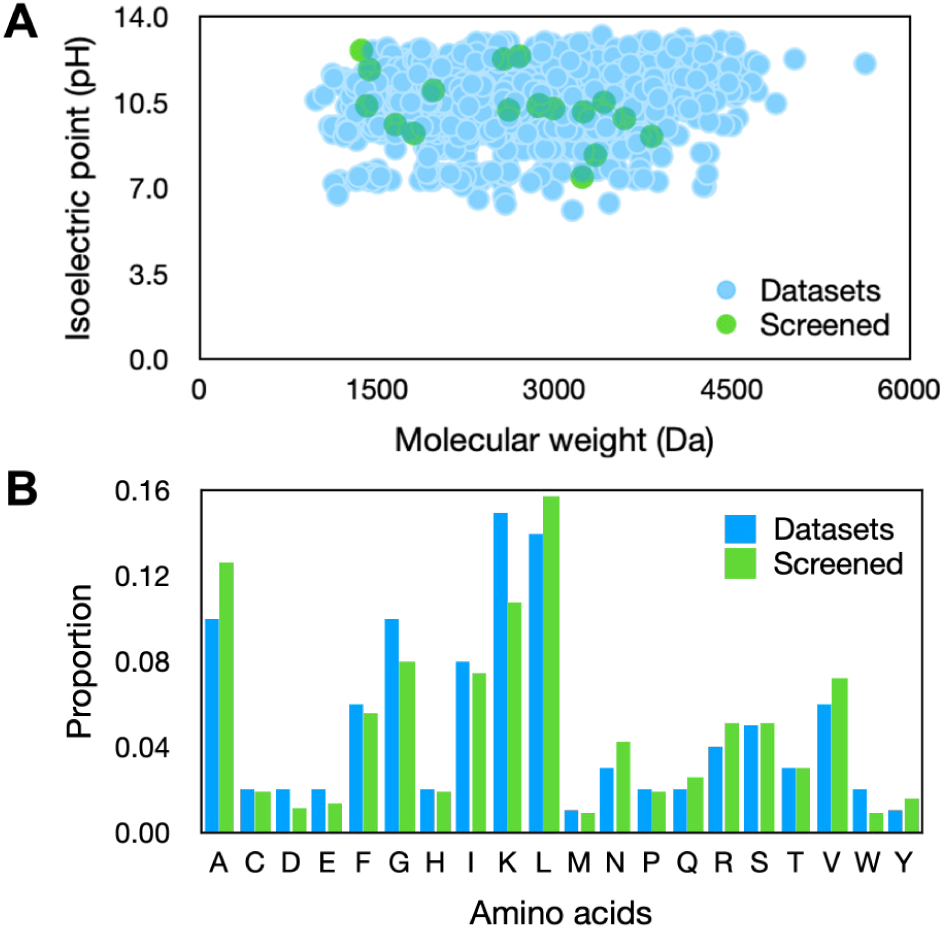
Charecteristic comparison between positive dataset and 20 screened peptides. (A) The distribution of isoelectric point and molecular weight. (B) Distribution of amino acids in the 20 screened peptides.

**Figure S2:**
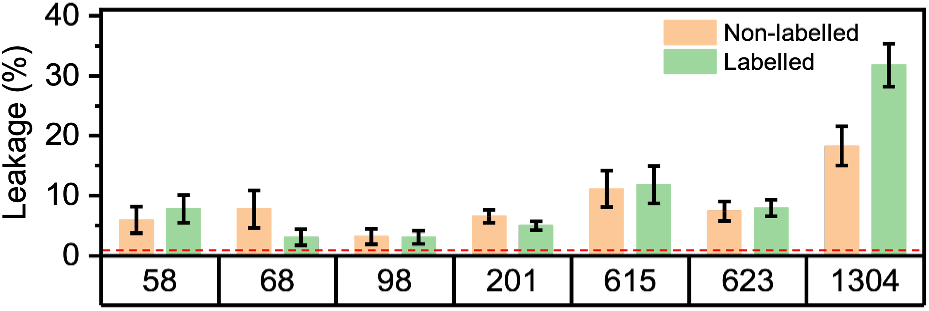
Leakage of carboxyfluorescein (CF) from liposomes induced by peptides and fluorophores-labeled peptides. Histogram represents the leakage of CF from liposomes after the addition of peptides, and is shown as a percentage of that induced by the addition of detergent 0.2% Triton X-100. Red dashed lines indicate the background leakage from the liposomes incubated with an equivalent concentration of buffer alone, to control for the effect of buffer. Data are presented as mean ± SEM (n = 3 independent measurements).

**Figure S3:**
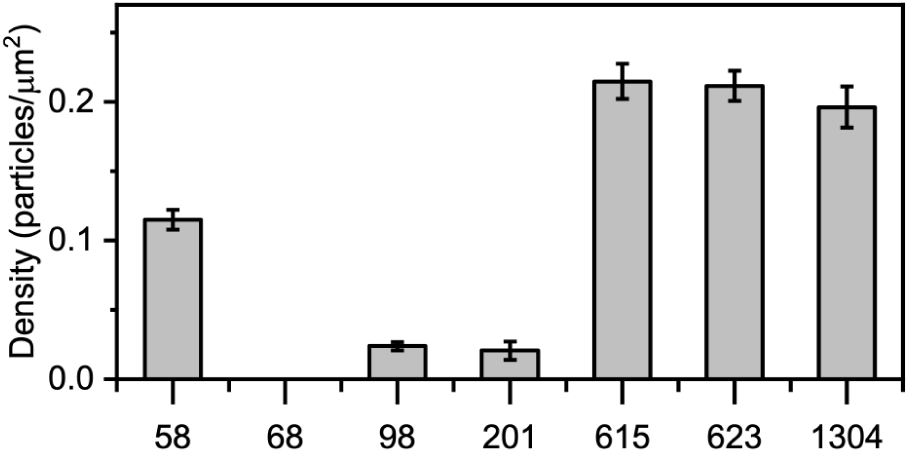
Surface density of peptides absorbing on lipid bilayer. Data are presented as mean ± SEM (n = 3 independent measurements).

**Figure S4:**
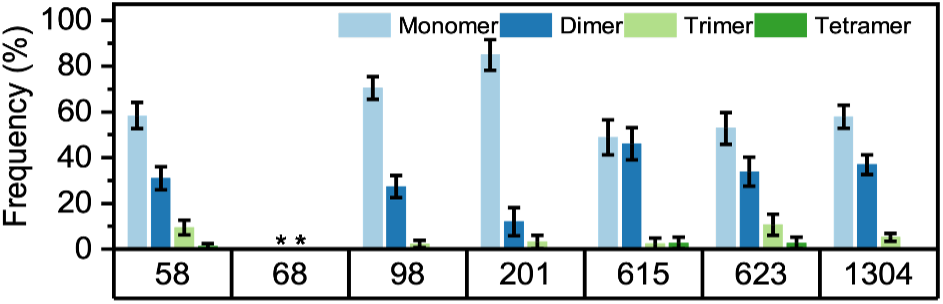
Oligomers distribution of peptides on supported lipid bilayers with washing free peptides in solution. Histogram of photobleaching step numbers for fluorescence labelled peptides analyzed. Asterisks represent the data of peptide 68 was not further analyzed. Data are presented as mean ± SD. Error bars correspond to the bootstrapped standard deviation.

**Figure S5:**
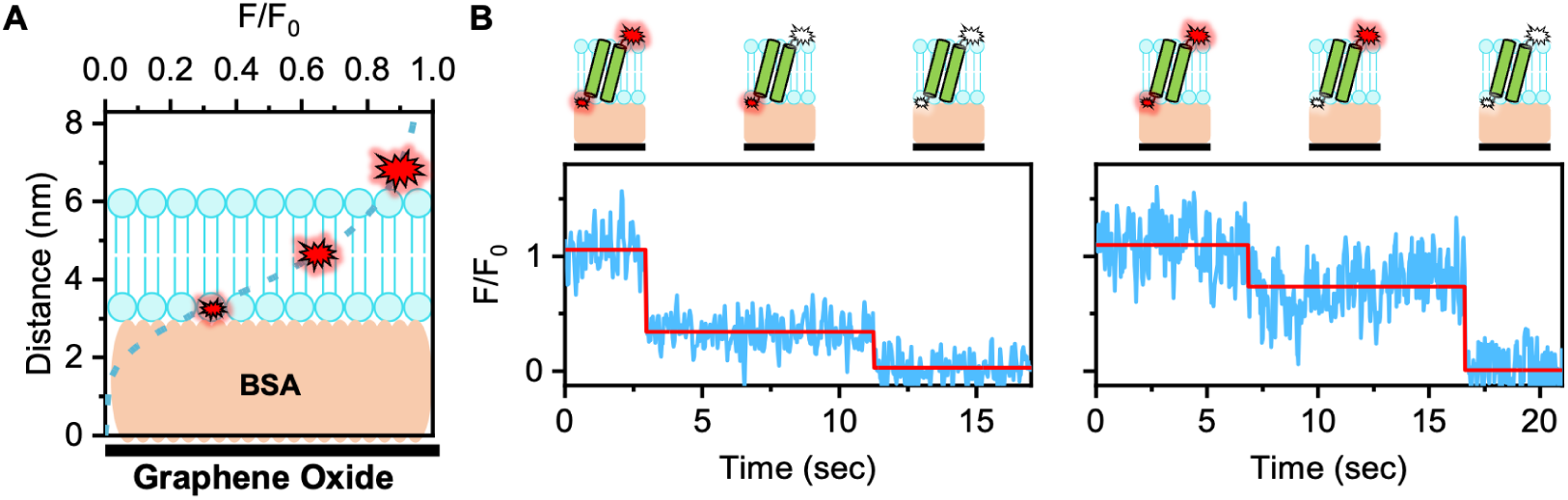
Detecting membrane topology of dimeric peptide 201 by smSIFA (single molecule surface-induced fluorescence attenuation). (A) The dependence of the intensity ratio on the distance to the surface of the single layer graphene oxide in sm-SIFA. (B) Two representative intensity ratio traces (intensity trace, blue; idealized fitting curves, red) of antiparallel dimeric peptide 201.

**Figure S6:**
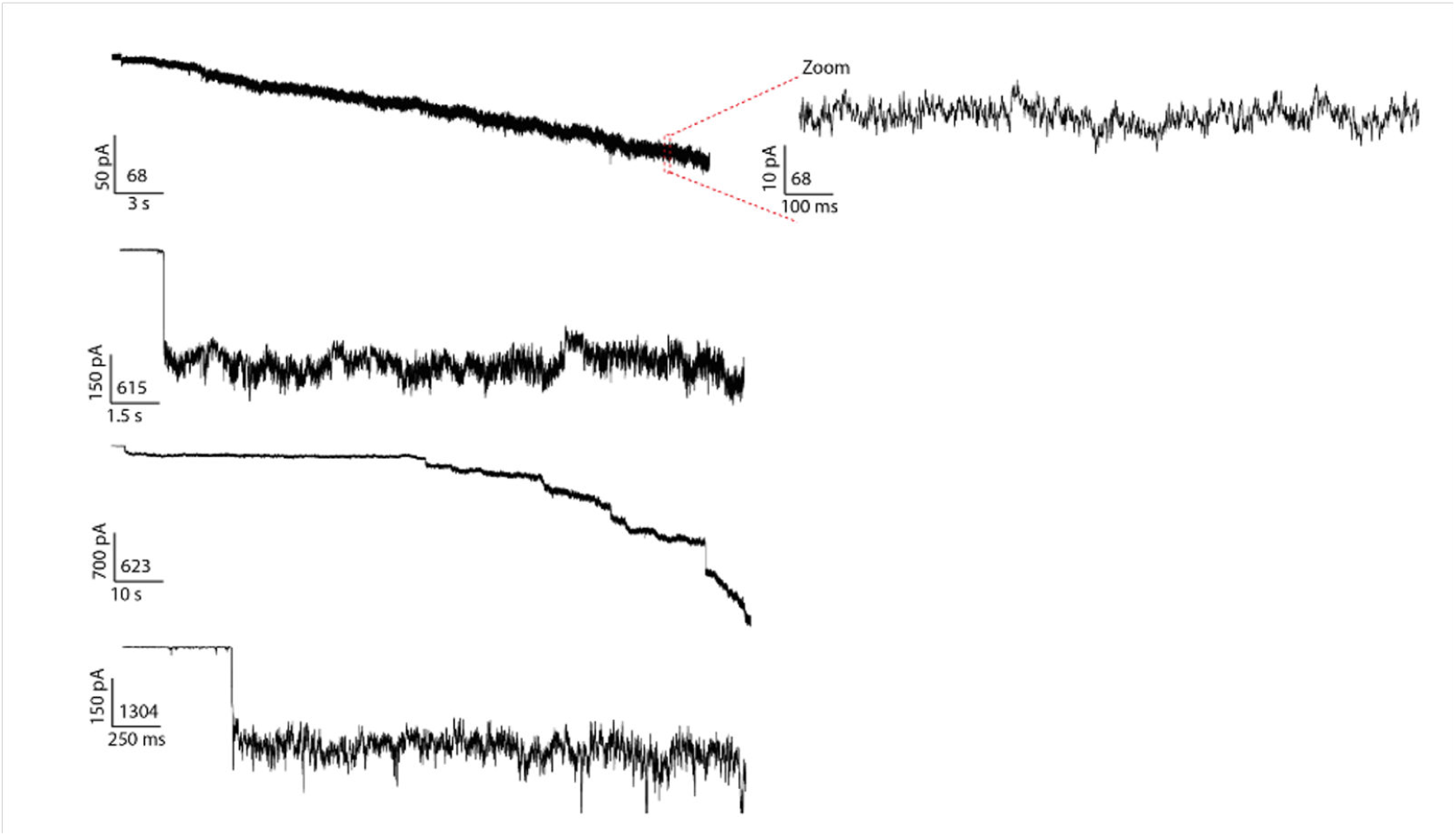
Single-channel bilayer experiments results for peptide 68, 615, 623, and 1304. These traces show disturbance of the peptide while in contact with the membrane. Peptides 615, 623 and 1304 show clear step indicative for pore insertion. For peptide 68, we do observe disturbance of the membrane with no clear steps like in the other peptides.

**Figure S7:**
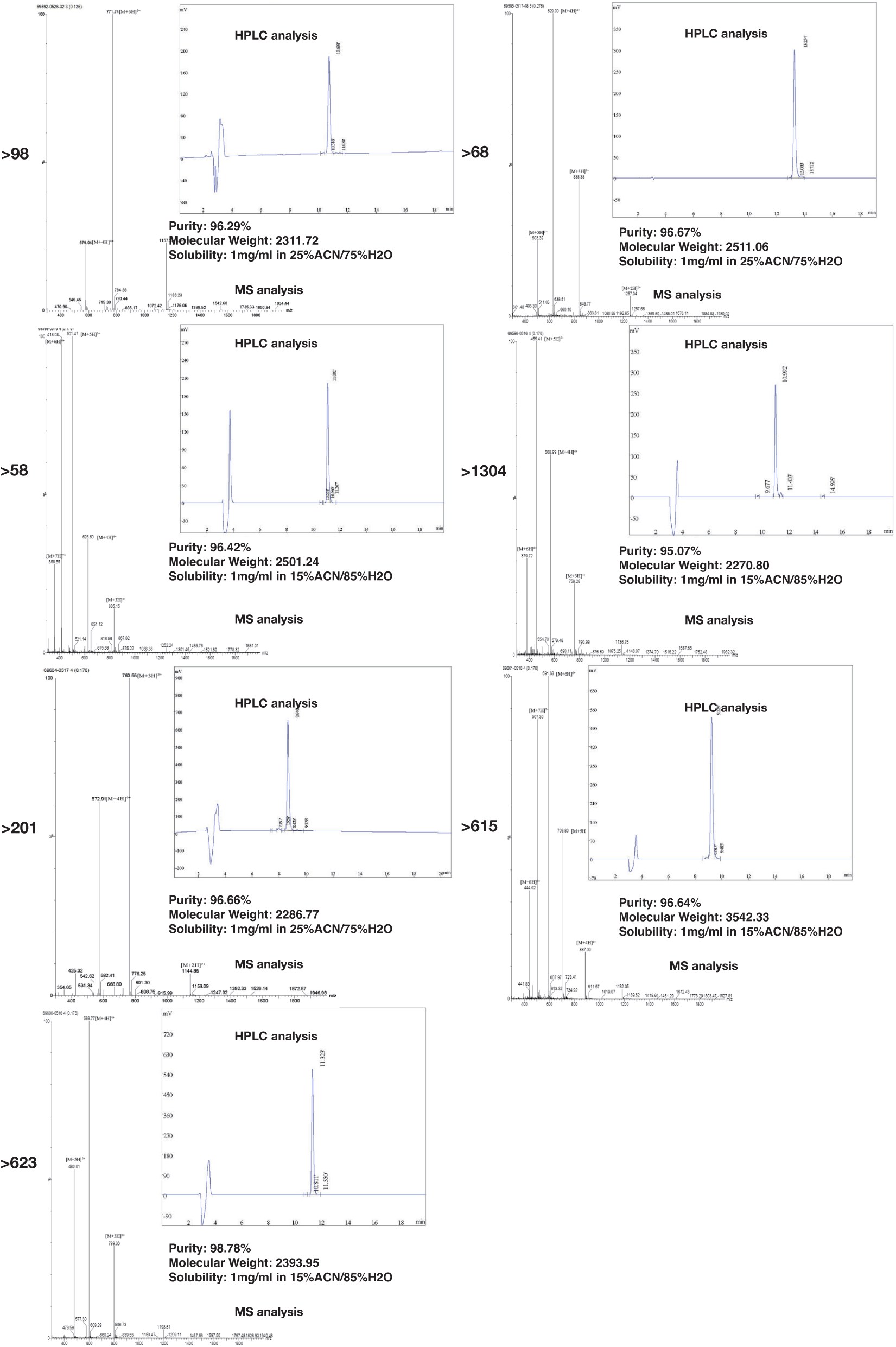
Synthesis reports of 7 effective peptides.

## References

[1] Lear, J. D.; Wasserman, Z. R.; DeGrado, W. F. Synthetic Amphiphilic Peptide Models for Protein Ion Channels. Science 1988, 240, 1177–1181.

[2] Mesa-Galloso, H.; Pedrera, L.; Ros, U. Pore-forming proteins: From defense factors to endogenous executors of cell death. Chemistry and Physics of Lipids 2021, 234, 105026.

[3] Guha, S.; Ghimire, J.; Wu, E.; Wimley, W. C. Mechanistic Landscape of Membrane-Permeabilizing Peptides. Chemical reviews 2019, 119, 6040—6085.

[4] Chen, N.; Jiang, C. Antimicrobial peptides: Structure, mechanism, and modification. European journal of medicinal chemistry 2023, 255, 115377.

[5] Yeaman, M.; Yount, N. Mechanisms of Antimicrobial Peptide Action and Resistance. Pharmacological reviews 2003, 55, 27–55.

[6] Mahlapuu, M.; Håkansson, J.; Ringstad, L.; Björn, C. Antimicrobial Peptides: An Emerging Category of Therapeutic Agents. Frontiers in Cellular and Infection Microbiology 2016, 6, 194.

[7] Yang, L.; Harroun, T.; Weiss, T.; Ding, L.; Huang, H. Barrel-Stave Model or Toroidal Model? A Case Study on Melittin Pores. Biophysical journal 2001, 81, 1475–85.

[8] Qian, S.; Wang, W.; Yang, L.; Huang, H. Structure of the Alamethicin Pore Reconstructed by X-Ray Diffraction Analysis. Biophysical journal 2008, 94, 3512–22.

[9] Song, C.; Weichbrodt, C.; Salnikov, E. S.; Dynowski, M.; Forsberg, B. O.; Bechinger, B.; Steinem, C.; de Groot, B. L.; Zachariae, U.; Zeth, K. Crystal structure and functional mechanism of a human antimicrobial membrane channel. Proceedings of the National Academy of Sciences 2013, 110, 4586–4591, Publisher: Proceedings of the National Academy of Sciences.

[10] Wiedman, G.; Fuselier, T.; He, J.; Searson, P. C.; Hristova, K.; Wimley, W. C. Highly Eicient Macromolecule-Sized Poration of Lipid Bilayers by a Synthetically Evolved Peptide. Journal of the American Chemical Society 2014, 136, 4724–4731, PMID: 24588399.

[11] Krauson, A. J.; Hall, O. M.; Fuselier, T.; Starr, C. G.; Kauffman, W. B.; Wimley, W. C. Conformational Fine-Tuning of Pore-Forming Peptide Potency and Selectivity. Journal of the American Chemical Society 2015, 137, 16144–16152, PMID: 26632653.

[12] Pillong, M.; Hiss, J. A.; Schneider, P.; Lin, Y.-C.; Posselt, G.; Pfeiffer, B.; Blatter, M.; Müller, A. T.; Bachler, S.; Neuhaus, C. S.; Dittrich, P. S.; Altmann, K.-H.; Wessler, S.; Schneider, G. Rational Design of Membrane-Pore-Forming Peptides. Small 2017, 13, 1701316.

[13] Mahendran, K. R.; Niitsu, A.; Kong, L.; Thomson, A. R.; Sessions, R. B.; Woolfson, D. N.; Bayley, H. A monodisperse transmembrane Ă-helical peptide barrel. Nature chemistry 2017, 9, 411–419.

[14] Krishnan R, S.; Jana, K.; Shaji, A.; Nair, K.; Das, A.; Vikraman, D.; Bajaj, H.; Kleinekathöfer, U.; Mahendran, K. Assembly of transmembrane pores from mirror-image peptides. Nature Communications 2022, 13, 5377.

[15] Chen, C. H.; Starr, C. G.; Troendle, E.; Wiedman, G.; Wimley, W. C.; Ulmschneider, J. P.; Ulmschneider, M. B. Simulation-Guided Rational de Novo Design of a Small Pore-Forming Antimicrobial Peptide. Journal of the American Chemical Society 2019, 141, 4839–4848.

[16] Ulmschneider, J.; Ulmschneider, M. Melittin can permeabilize membranes via large transient pores. Nature Communications 2024, 15, 7281.

[17] Deb, R. et al. Computational Design of Pore-Forming Peptides with Potent An-timicrobial and Anticancer Activities. Journal of Medicinal Chemistry 2024, 67, 14040–14061.

[18] Ma, Y.; Guo, Z.; Xia, B.; Zhang, Y.; Liu, X.; Yu, Y.; Tang, N.; Tong, X.; Wang, M.; Ye, X.; Feng, J.; Chen, Y.; Wang, J. Identification of antimicrobial peptides from the human gut microbiome using deep learning. Nature Biotechnology 2022, 40, 1–11.

[19] Santos-Júnior, C. D. et al. Discovery of antimicrobial peptides in the global microbiome with machine learning. Cell 2024, 187, 3761–3778.e16.

[20] Torres, M.; Brooks, E.; Cesaro, A.; Sberro, H.; Gill, M.; Nicolaou, C.; Bhatt, A.; de la Fuente, C. Mining human microbiomes reveals an untapped source of peptide antibiotics. Cell 2024, 187, 5453–5467.e15.

[21] Torres, M. D.; Melo, M. C.; Crescenzi, O.; Notomista, E.; de la Fuente-Nunez, C. Mining for encrypted peptide antibiotics in the human proteome. Nature Biomedical Engineering 2022, 6, 67–75.

[22] Maasch, J.; Torres, M.; Cardoso dos Reis Melo, M.; de la Fuente, C. Molecular deextinction of ancient antimicrobial peptides enabled by machine learning. Cell Host & Microbe 2023, 31, 1260–1274.e6.

[23] Wan, F.; Torres, M.; Peng, J.; de la Fuente, C. Deep-learning-enabled antibiotic discovery through molecular de-extinction. Nature Biomedical Engineering 2024, 8, 1–18.

[24] Huang, J.; Xu, Y.; Xue, Y.; Huang, Y.; Li, X.; Chen, X.; Xu, Y.; Zhang, D.; Zhang, P.; Zhao, J.; Ji, J. Identification of potent antimicrobial peptides via a machine-learning pipeline that mines the entire space of peptide sequences. Nature Biomedical Engineering 2023, 7, 1–14.

[25] Wang, L.; Zhang, J.; Wang, D.; Song, C. Membrane contact probability: An essential and predictive character for the structural and functional studies of membrane proteins. PLoS Computational Biology 2022, 18, e1009972.

[26] Elnaggar, A.; Heinzinger, M.; Dallago, C.; Rehawi, G.; Wang, Y.; Jones, L.; Gibbs, T.; Feher, T.; Angerer, C.; Steinegger, M.; Bhowmik, D.; Rost, B. ProtTrans: towards cracking the language of life’s code through self-supervised deep learning and high performance computing. CoRR 2020,

[27] Unsal, S.; Atas, H.; Albayrak, M.; Turhan, K.; Acar, A. C.; Doğan, T. Learning functional properties of proteins with language models. *Nat*. Mach. Intell. 2022, 4, 227–245.

[28] Wang, G.; Li, X.; Wang, Z. APD3: the antimicrobial peptide database as a tool for research and education. Nucleic Acids Res. 2016, 44, D1087–D1093.

[29] Pirtskhalava, M.; Amstrong, A. A.; Grigolava, M.; Chubinidze, M.; Alimbarashvili, E.; Vishnepolsky, B.; Gabrielian, A.; Rosenthal, A.; Hurt, D. E.; Tartakovsky, M. DBAASP v3: database of antimicrobial/cytotoxic activity and structure of peptides as a resource for development of new therapeutics. Nucleic Acids Research 2020, 49, D288–D297.

[30] Piotto, S. P.; Sessa, L.; Concilio, S.; Iannelli, P. YADAMP: yet another database of antimicrobial peptides. International Journal of Antimicrobial Agents 2012, 39, 346–351.

[31] Singh, S.; Chaudhary, K.; Dhanda, S. K.; Bhalla, S.; Usmani, S. S.; Gautam, A.; Tuknait, A.; Agrawal, P.; Mathur, D.; Raghava, G. P. SATPdb: A database of structurally annotated therapeutic peptides. Nucleic Acids Research 2015, 44, D1119–D1126.

[32] Suzek, B. E.; Wang, Y.; Huang, H.; McGarvey, P. B.; Wu, C. H.; the UniProt Consortium UniRef clusters: a comprehensive and scalable alternative for improving sequence similarity searches. Bioinformatics 2014, 31, 926–932.

[33] Lee, H.; Lee, S.; Lee, I.; Nam, H. AMP-BERT: Prediction of antimicrobial peptide function based on a BERT model. Protein Science 2023, 32.

[34] Lawrence, T. J.; Carper, D. L.; Spangler, M. K.; Carrell, A. A.; Rush, T. A.; Minter, S. J.; Weston, D. J.; Labbé, J. L. amPEPpy 1.0: a portable and accurate antimicrobial peptide prediction tool. Bioinformatics 2020, 37, 2058–2060.

[35] Li, C.; Sutherland, D.; Hammond, S. A.; Yang, C.; Taho, F.; Bergman, L.; Houston, S.; Warren, R. L.; Wong, T.; Hoang, L. M.; Cameron, C. E.; Helbing, C. C.; Birol, I. AMPlify: attentive deep learning model for discovery of novel antimicrobial peptides effective against WHO priority pathogens. BMC Genomics 2022, 23.

[36] Santos-Júnior, C. D.; Pan, S.; Zhao, X. M.; Coelho, L. P. Macrel: Antimicrobial peptide screening in genomes and metagenomes. PeerJ 2020, 8.

[37] Veltri, D.; Kamath, U.; Shehu, A. Deep learning improves antimicrobial peptide recognition. Bioinformatics 2018, 34, 2740–2747.

[38] Cao, Q. et al. Designing antimicrobial peptides using deep learning and molecular dynamic simulations. Briefings in Bioinformatics 2023,

[39] Li, J.; Wang, L.; Zhu, Z.; Song, C. Exploring the Alternative Conformation of a Known Protein Structure Based on Contact Map Prediction. Journal of Chemical Information and Modeling 2024, 64.

[40] Song, C.; de Groot, B. L.; Sansom, M. S. Lipid Bilayer Composition Influences the Activity of the Antimicrobial Peptide Dermcidin Channel. Biophysical Journal 2019, 116, 1658–1666.

[41] Perez-Riverol, Y.; Bai, J.; Bandla, C.; García-Seisdedos, D.; Hewapathirana, S.; Kamatchinathan, S.; Kundu, D.; Prakash, A.; Frericks-Zipper, A.; Eisenacher, M.; Walzer, M.; Wang, S.; Brazma, A.; Vizcaíno, J. The PRIDE database resources in 2022: a hub for mass spectrometry-based proteomics evidences. Nucleic Acids Research 2021, 50, D543–D552.

[42] Zhang, H.; Guo, P. Single molecule photobleaching (SMPB) technology for counting of RNA, DNA, protein and other molecules in nanoparticles and biological complexes by TIRF instrumentation. Methods 2014, 67, 169–176, Nucleic Acids Nanotechnology.

[43] Jiang, H.; Chen, W.; Wang, J.; Zhang, R. Selective N-terminal modification of peptides and proteins: Recent progresses and applications. Chinese Chemical Letters 2022, 33, 80–88.

[44] Li, Y.; Qian, Z.; Ma, L.; Hu, S.; Nong, D.; Xu, C.; Ye, F.; Lu, Y.; Wei, G.; Li, M. Single-molecule visualization of dynamic transitions of pore-forming peptides among multiple transmembrane positions. Nature Communications 2016, 7.

[45] Yang, C.; Ma, D.; Hu, S.; Li, M.; Lu, Y. Real-time analysis of nanoscale dynamics in membrane protein insertion via single-molecule imaging. Biophysics Reports 2024, 10, 1–8.

[46] Huan, Y.; Kong, Q.; Mou, H.; Yi, H. Antimicrobial Peptides: Classification, Design, Application and Research Progress in Multiple Fields. Frontiers in Microbiology 2020, 11.

[47] Szymczak, P.; Możejko, M.; Grzegorzek, T.; Jurczak, R.; Bauer, M.; Neubauer, D.; Sikora, K.; Michalski, M.; Sroka, J.; Setny, P.; Kamysz, W.; Szczurek, E. Discovering highly potent antimicrobial peptides with deep generative model HydrAMP. Nature communications 2023, 14, 1453.

[48] Verma, D. P.; Tripathi, A. K.; Thakur, A. K. Innovative Strategies and Methodologies in Antimicrobial Peptide Design. Journal of Functional Biomaterials 2024, 15.

[49] Liu, H.; Song, Z.; Zhang, Y.; Wu, B.; Chen, D.; Zhou, Z.; Zhang, H.; Li, S.; Feng, X.; Huang, J.; Wang, H. De novo design of self-assembling peptides with antimicrobial activity guided by deep learning. Nature Materials 2025,

[50] Heffernan, R.; Paliwal, K.; Lyons, J.; Singh, J.; Yang, Y.; Zhou, Y. Single-sequence-based prediction of protein secondary structures and solvent accessibility by deep whole-sequence learning. Journal of Computational Chemistry 2018, 39, 2210–2216.

[51] Kabsch, W.; Sander, C. Dictionary of protein secondary structure: pattern recognition of hydrogen-bonded and geometrical features. Biopolymers: Original Research on Biomolecules 1983, 22, 2577–2637.

[52] Wen, Z.; Shi, J.; Li, Q.; He, B.; Chen, J. ThunderSVM: a fast SVM library on GPUs and CPUs. J. Mach. Learn. Res. 2018, 19, 797–801.

[53] Camacho, C.; Coulouris, G.; Avagyan, V.; Ma, N.; Papadopoulos, J.; Bealer, K.; Madden, T. L. BLAST+: architecture and applications. BMC Bioinformatics 2009, 10, 421.

[54] Das, P.; Sercu, T.; Wadhawan, K.; Padhi, I.; Gehrmann, S.; Cipcigan, F.; Chenthamarakshan, V.; Strobelt, H.; dos Santos, C.; Chen, P. Y.; Yang, Y. Y.; Tan, J. P.; Hedrick, J.; Crain, J.; Mojsilovic, A. Accelerated antimicrobial discovery via deep generative models and molecular dynamics simulations. Nature Biomedical Engineering 2021, 5, 613–623.

[55] Müller, A. T.; Gabernet, G.; Hiss, J. A.; Schneider, G. modlAMP: Python for antimicrobial peptides. Bioinformatics 2017, 33, 2753–2755.

[56] Kyte, J.; Doolittle, R. F. A simple method for displaying the hydropathic character of a protein. Journal of Molecular Biology 1982, 157, 105–132.

[57] Zeng, H.; Wang, S.; Zhou, T.; Zhao, F.; Li, X.; Wu, Q.; Xu, J. ComplexContact: a web server for inter-protein contact prediction using deep learning. Nucleic acids research 2018, 46, W432–W437.

[58] Zhao, J.; Pei, S.; Ren, W.; Gao, L.; Cheng, H. M. Eicient preparation of largearea graphene oxide sheets for transparent conductive films. ACS Nano 2010, 4, 5245–5252.

[59] Shi, L.; Yang, C.; Zhang, M.; Li, K.; Wang, K.; Jiao, L.; Liu, R.; Wang, Y.; Li, M.; Wang, Y.; Ma, L.; Hu, S.; Bian, X. Dissecting the mechanism of atlastin-mediated homotypic membrane fusion at the single-molecule level. Nature Communications 2024, 15.

## References

[1] Veltri, D.; Kamath, U.; Shehu, A. Deep learning improves antimicrobial peptide recognition. Bioinformatics 2018, 34, 2740–2747.

[2] Santos-Júnior, C. D.; Pan, S.; Zhao, X. M.; Coelho, L. P. Macrel: Antimicrobial peptide screening in genomes and metagenomes. PeerJ 2020, 8.

[3] Lawrence, T. J.; Carper, D. L.; Spangler, M. K.; Carrell, A. A.; Rush, T. A.; Minter, S. J.; Weston, D. J.; Labbé, J. L. amPEPpy 1.0: a portable and accurate antimicrobial peptide prediction tool. Bioinformatics 2020, 37, 2058–2060.

[4] Li, C.; Sutherland, D.; Hammond, S. A.; Yang, C.; Taho, F.; Bergman, L.; Houston, S.; Warren, R. L.; Wong, T.; Hoang, L. M.; Cameron, C. E.; Helbing, C. C.; Birol, I. AM-Plify: attentive deep learning model for discovery of novel antimicrobial peptides effective against WHO priority pathogens. BMC Genomics 2022, 23.

[5] Lee, H.; Lee, S.; Lee, I.; Nam, H. AMP-BERT: Prediction of antimicrobial peptide function based on a BERT model. Protein Science 2023, 32.

[6] Cao, Q. et al. Designing antimicrobial peptides using deep learning and molecular dynamic simulations. Briefings in Bioinformatics 2023,

[7] Li, C.; Zou, Q.; Jia, C.; Zheng, J. AMPpred-MFA: An Interpretable Antimicrobial Peptide Predictor with a Stacking Architecture, Multiple Features, and Multihead Attention. Journal of Chemical Information and Modeling 2024, 64, 2393–2404.

